# Chromosome-scale genomes of two wild flowering cherrys (*Cerasus itosakura* and *C. jamasakura*) provide insights into structural evolution in *Prunus*

**DOI:** 10.1101/2025.08.30.673203

**Authors:** Kazumichi Fujiwara, Atsushi Toyoda, Toshio Katsuki, Yutaka Sato, Bhim B. Biswa, Takushi Kishida, Momi Tsuruta, Yasukazu Nakamura, Takako Mochizuki, Noriko Kimura, Shoko Kawamoto, Tazro Ohta, Ken-Ichi Nonomura, Hironori Niki, Hiroyuki Yano, Kinji Umehara, Chikahiko Suzuki, Tsuyoshi Koide

**Author notes:** Author for correspondence: Tsuyoshi Koide, Mouse Genomics Resource Laboratory, National Institute of Genetics 1111 Yata, Mishima, Shizuoka, 411-8540, Japan.

## Abstract

Flowering cherries (genus *Cerasus*) are iconic trees in Japan, celebrated for their cultural and ecological significance. Despite their prominence, high-quality genomic resources for wild *Cerasus* species have been limited. Here, we report chromosome-level genome assemblies of two representative Japanese cherries: *Cerasus itosakura*, a progenitor of the widely cultivated *C.* ×*yedoensis* ‘Somei-yoshino’, and *Cerasus jamasakura*, a traditional popular wild species endemic to Japan. Using deep PacBio long-read and Illumina short-read sequencing, combined with reference-guided scaffolding based on near-complete *C. speciosa* genome, we generated assemblies of 259.1 Mbp (*C. itosakura*) and 312.6 Mbp (*C. jamasakura*), with both >98% BUSCO completeness. Consistent with their natural histories, *C. itosakura* showed low heterozygosity, while *C. jamasakura* displayed high genomic diversity. Comparative genomic analyses revealed structural variations, including large chromosomal inversions. Notably, the availability of both the previously published *C. speciosa* genome and our new *C. itosakura* genome enabled the reconstruction of proxy haplotypes for both parental lineages of ‘Somei-yoshino’. Comparison with the phased genome of ‘Somei-yoshino’ revealed genomic discrepancies, suggesting that the cultivar may have arisen from genetically distinct or admixed individuals, and may also reflect intraspecific diversity. Our results offer genomic foundations for evolutionary and breeding studies in *Cerasus* and *Prunus*.

## Introduction

Flowering cherries (genus *Cerasus*, formerly classified under *Prunus* subgenus *Cerasus*) represent a distinct lineage of deciduous trees indigenous to East Asia. In Japan, they hold exceptional cultural and historical significance. The custom of cherry blossom viewing, or *hanami*, dates back over a millennium and continues to inspire literature, visual art, and seasonal festivities^1–3^. This deep cultural association has driven long-standing horticultural innovation, including selective breeding, exploitation of spontaneous bud mutations, and interspecific hybridization, ultimately producing an extraordinary range of ornamental cultivars. Among these, the Sato-zakura Group^4^, which is predominantly derived from *Cerasus speciosa*, comprises many of the traditional cultivars developed in Japan^5^.

The Japanese archipelago harbors approximately ten native wild *Cerasus* species, which occupy a wide spectrum of ecological habitats, including coastal islands, riparian zones, and subalpine forests^6–10^. These species display substantial variation in floral morphology, fragrance composition, phenology, and growth habit. Three species, in particular, have played central roles in the cultural and horticultural history of Japanese cherries: *C. itosakura*, *C. jamasakura*, and *C. speciosa*. *C. itosakura* is a species in which flowers emerge before the leaves, by pale pink to white petals and exceptional longevity, with some individuals reportedly surviving for centuries^2,6,8,11^. These attributes have made it a preferred species for symbolic plantings at religious and cultural sites, and it has served as a progenitor in the development of several ornamental cultivars. *C. jamasakura*, in contrast, is a hill-dwelling species that typically flowers alongside leaf emergence, and has historically been the most familiar cherry in satoyama regions, where human settlements and low hills intersect. ^2,6,8,11^. The visual harmony of its flowers and young foliage has been celebrated in classical Japanese literature and art as a quintessential representation of spring. *C. speciosa*, native to maritime environments, produces large, fragrant flowers rich in coumarins and displays adaptations to coastal conditions^2,6,8,11^. This species has contributed substantially to the origin of many cultivars within the Sato-zakura Group.

These three species represent the principal genetic foundation of modern ornamental cherries. A prominent example is *C.* ×*yedoensis* ‘Somei-yoshino’, a clonally propagated interspecific hybrid derived from *C. itosakura* and *C. speciosa*^12–15^. This cultivar integrates the early-flowering trait of *C. itosakura* with the floral size of *C. speciosa*, resulting in a highly synchronous and visually striking floral display. Today, ‘Somei-yoshino’ is the most extensively planted cherry cultivar in Japan and has been widely introduced to temperate regions worldwide, where it serves as the focal point of public hanami festivals.

Advances in genome sequencing have greatly enhanced our understanding of cherry evolution and domestication. The near-complete genome of *C. speciosa* resolved centromeric repeat structures and provided a chromosome-scale, gap-free diploid reference^16^. Likewise, the assembly of *C. campanulata* ‘Lianmeiren’ achieved near-complete continuity and uncovered widespread structural variation, including single-nucleotide polymorphisms, small indels, and large-scale rearrangements^17^. These genomic resources have significantly expanded the framework for comparative analyses within *Cerasus*.

Despite these developments, chromosome-scale assemblies have not been available for *C. itosakura* or *C. jamasakura*, limiting systematic genomic comparisons and preventing detailed assessment of structural variation and haplotype composition in hybrid cultivars such as ‘Somei-yoshino’^18^. The long-read assembly of this hybrid remains unanchored to chromosomes, limiting its utility in elucidating parental contributions.

Recent phylogenomic and morphological studies have also led to a revision of the taxonomic status of flowering cherries. *Cerasus* was previously classified as a subgenus of *Prunus*. However, recent studies based on nuclear and chloroplast genome sequences, together with comparative morphological data, support its recognition as an independent genus^11,19–21^. Plastid-based phylogenies reinforce its monophyly and highlight its deep evolutionary divergence from other *Prunus* lineages, underscoring the distinct evolutionary trajectory of the group.

In this study, we present high-quality, chromosome-scale genome assemblies for *C. itosakura* and *C. jamasakura*. These assemblies were generated using a reference-guided scaffolding approach based on the near-complete genome of *C. speciosa*. We additionally assembled the complete chloroplast and mitochondrial genomes for both species. Structural annotation included the identification of protein-coding genes, repetitive elements, and non-coding RNAs. To investigate the genomic context of interspecific hybridization, we reconstructed the previously published contig-level assembly of ‘Somei-yoshino’ and compared it to the new parental genomes.

The addition of chromosome-level assemblies for *C. itosakura* and *C. jamasakura*, alongside existing high-contiguity genomes for *C. speciosa* and *C. campanulata*, establishes a comprehensive genomic framework for the genus *Cerasus*. This resource enables fine-resolution analyses of genome architecture, synteny, hybridization, and the genetic basis of key ornamental traits, including flowering time, floral morphology, fragrance biosynthesis, and stress responses. These genomic references also provide essential tools for evolutionary biology, conservation genomics, and molecular breeding in one of the most culturally significant groups of flowering plants.

## Materials and methods

### Plant materials

The young leaf buds of *Cerasus itosakura* and *Cerasus jamasakura* var. *jamasakura* were collected from the following sources: for *C. itosakura*, samples were obtained from a grafted individual (NIG-0593) propagated from a naturally growing tree designated as a natural monument of Izu City, located at Sugimoto-bashi in Amagi, Shizuoka Prefecture, Japan (Fig. 1A, B); for *C. jamasakura*, samples were directly collected from a wild individual (TFSG-CP70p1p4) growing in the Tama Forest Science Garden of the Forestry and Forest Products Research Institute, located in Hachioji City, Tokyo, Japan (Fig. 1C, D).

**Fig. 1.**
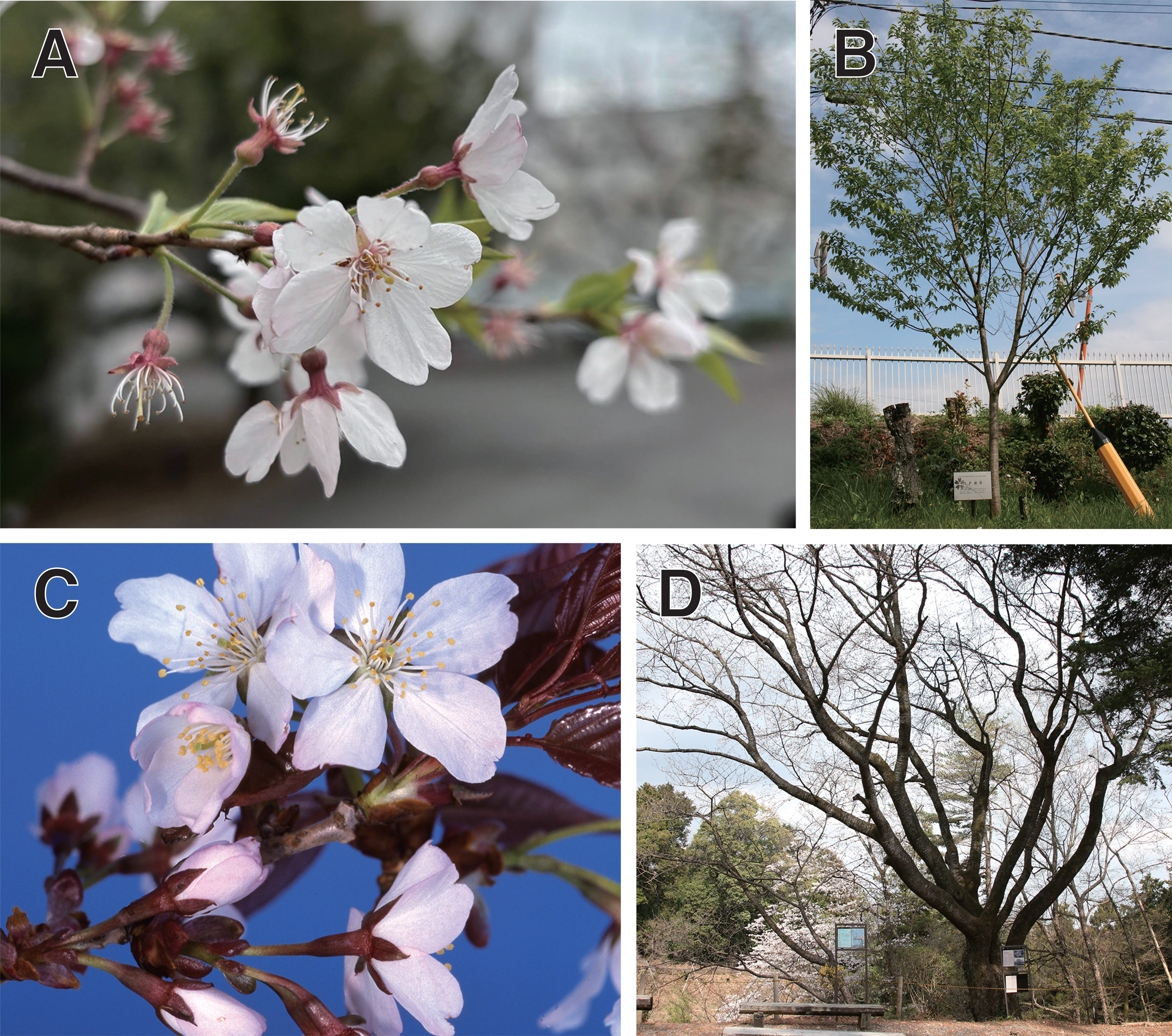
Photographs of the two *Cerasus* species examined in this study. (A) Close-up of *C. itosakura* flowers. (B) Entire tree of *C. itosakura*. (C) Close-up of *C. jamasakura* flowers. (D) Entire tree of *C. jamasakura*. Panels C and D were provided by the Tama Forest Science Garden, Forestry and Forest Products Research Institute.

### Sampling collection

Leaves were harvested, rinsed with deionized water, gently blotted dry using paper towels, and immediately stored at −80 °C until genomic DNA extraction. DNA was isolated using a modified version of a previously described protocol. In brief, 0.1– 0.5 g of leaf tissue was flash-frozen in liquid nitrogen and ground to a fine powder with a pre-chilled mortar and pestle. The powder was suspended in 10 mL of Carlson lysis buffer (100 mM Tris-HCl, pH 9.0, 2% CTAB, 1.4 M NaCl, 1% PEG 6000, 1% polyvinylpyrrolidone K30, and 20 mM EDTA), supplemented with 0.25% β-mercaptoethanol. The lysate was incubated at 74 °C for 20 minutes with gentle inversion every 5 minutes. After cooling to room temperature, 20 mL of chloroform:isoamyl alcohol (24:1) was added, mixed thoroughly, and centrifuged at 1,800 × g for 10 minutes. The aqueous phase was transferred to a fresh tube, mixed with an equal volume of 2-propanol, and centrifuged at 2,500 × g for 30 minutes. The resulting pellet was washed with 75% ethanol and centrifuged again for 10 minutes. After discarding the supernatant, the pellet was briefly air-dried and resuspended in 1 mL of TE buffer. RNase was added to a final concentration of 10–100 µg/mL, and the sample was incubated overnight at 4 °C. Genomic DNA was further purified using the QIAGEN Genomic-tip 500 G and DNA Buffer Set (QIAGEN, Hilden, Germany), following the manufacturer’s instructions.

### Library preparation and genome sequencing

For Illumina sequencing, genomic DNA was sheared using a Covaris system and libraries were prepared with the TruSeq DNA PCR-Free Library Prep Kit (Illumina, CA, USA). Size selection was carried out by agarose gel excision, and 250 bp paired-end sequencing was performed on the NovaSeq 6000 platform using a 500-cycle NovaSeq6000 Reagent Kit (Illumina, CA, USA). For continuous long-read (CLR) sequencing on the PacBio Sequel II platform, DNA was fragmented using a g-TUBE, and libraries were constructed with the SMRTbell Express Template Prep Kit 2.0 (Pacific Biosciences, CA, USA). DNA fragments smaller than 40 kb were excluded using BluePippin, and sequencing was conducted with the Sequel II Binding Kit 2.0 and Sequencing Kit 2.0 (Pacific Biosciences, CA, USA).

### Genome size estimation

The genome sizes of *C. itosakura* and *C. jamasakura* were estimated using a *k*-mer– based approach with Illumina short-read sequencing data. Raw reads underwent stringent quality filtering using fastp (v0.23.2)^22^ with the parameters -q 30 -f 5 -F 5 -t 5 - T 5 -n 0 -u 20. Canonical 21-mers were then counted using KMC3 (v3.2.1)^23^ with the settings -k 21 -ci 1 -cs 10000. The *k*-mer size of 21 was selected following established guidelines for genome size estimation, as it strikes a balance between minimizing repetitive *k*-mers and maintaining robustness against sequencing errors. K-mer frequency histograms were generated using kmc_tools with the operations transform and histogram (-cx 10000). These histograms were subsequently analyzed with GenomeScope 2.0 (Rscript version, v2.0)^24,25^ using the parameters -k 21 -p 2 to estimate haploid genome size and heterozygosity. Given that most cherry blossom species in Japan, including *C. itosakura* and *C. jamasakura*, are cytologically confirmed to be diploid, a diploid model (-p 2) was applied in GenomeScope analyses.

### Genome assembly

The primary genome assemblies of *C. itosakura* and *C. jamasakura* were generated from long-read sequencing data using the Canu (v2.2)^26^ assembler. Canu was executed with the following parameters: genomeSize=250m, corOutCoverage=60, correctedErrorRate=0.045, corMaxEvidenceErate=0.15, batOptions=“-dg 3 -db 3 -dr 1 - ca 500 -cp 50”, and -fast. These settings were selected based on recommendations in the official Canu documentation. During post-assembly processing, contigs flagged with the tag “SuggestBubble=yes” were excluded to eliminate potentially spurious sequences.

To screen the raw, bubble-free contigs for potential contamination, we used blastn from the NCBI BLAST+ suite (v2.12.0+)^27^ against the NCBI Prokaryotic RefSeq Genomes database, applying an E-value threshold of “1e-10”. Contaminant sequences were identified and removed using BlobTools (v1.0)^28^, which integrates sequence composition and taxonomic information to flag non-target contigs.

Each filtered raw assembly was then polished using its corresponding long-read data. Long reads were first aligned to the draft assemblies using minimap2^29^ implemented in pbmm2 (v1.16.0) (https://github.com/PacificBiosciences/pbmm2), and polishing was performed with the Arrow algorithm implemented in GCpp (v2.0.2) (https://github.com/PacificBiosciences/gcpp). This polishing step was iterated twice to further improve consensus quality.

Following polishing, we performed haplotig purging to remove redundant contigs originating from heterozygous regions. To address this, we employed Purge_haplotigs (v1.1.3)^30^, which distinguishes primary contigs from haplotigs using a combination of coverage depth and repeat annotation.

To generate high-confidence repeat annotations, we first conducted de novo repeat identification using RepeatModeler2 (v2.0.5)^31^ with the -LTRStruct option enabled, facilitating detection of long terminal repeat (LTR) retrotransposons. The resulting repeat library was then used in RepeatMasker (v4.1.5)^32^ to annotate repetitive regions. These annotations were supplied to Purge_haplotigs via the -r option to assist haplotig classification. The tool was run with default parameters, except for customized coverage thresholds: -l 50 -m 370 -h 700 for *C. itosakura*, and -l 50 -m 290 -h 550 for *C. jamasakura*.

The purged primary assemblies were further refined by removing chloroplast and mitochondrial sequences. To identify organellar contigs, we aligned the assemblies to the complete chloroplast and mitochondrial genomes of *C. itosakura* and *C. jamasakura* (described below) using nucmer from MUMmer4 (v4.0.0rc1)^33^. Alignments were visualized and manually inspected to identify and remove contigs of organellar origin based on coverage and sequence similarity.

Finally, the purged assemblies, free of organellar sequences, were subjected to short-read polishing. We used NextPolish (v1.4.1)^34^ with default settings for two rounds of short-read-based polishing to enhance sequence accuracy for downstream analyses.

### Reference-guided scaffolding

High-quality, contig-level assemblies of *C. itosakura* and *C. jamasakura* were further scaffolded to chromosome-level assemblies using RagTag (v2.1.0)^35^, a reference-guided scaffolding tool. For this process, we used the complete genome assembly of *Cerasus speciosa* as the reference. The use of *C. speciosa* as a reference for RagTag scaffolding is justified by its close phylogenetic relationship to both *C. itosakura* and *C. jamasakura*, all of which belong to the genus *Cerasus*. These species share a conserved karyotype (2n = 2x = 16)^36–39^ and exhibit substantial genome collinearity, as shown in previous comparative genomic studies of *Prunus* and *Cerasus* species. The overall workflow from genome assembly to reference-guided scaffolding is summarized in Supplementary Fig. 1.

### Chloroplast and Mitochondria assembly

The chloroplast genomes of *C. itosakura* and *C. jamasakura* were de novo assembled from Illumina short-read data. Reads were quality-filtered and trimmed using fastp, as described above. Clean reads were then assembled using the get_organelle_from_reads.py script from GetOrganelle (v1.7.7.0)^40^ with the parameters -R 15 -k 21,55,85,115,127 -F embplant_pt. Among the resulting assemblies, the one generated with a *k*-mer size of 127 was selected for downstream analysis. Gene annotation of the assembled circular chloroplast genomes was performed using GeSeq^41^ (https://chlorobox.mpimp-golm.mpg.de/geseq.html). While default settings were largely retained, the following adjustments were made: the input FASTA file was specified as circular, the sequence source was set to “plastid (land plants),” and Cerasus speciosa (NCBI RefSeq: NC_043921.1) was used as the reference for BLAT searches. Additionally, Chloë v0.1.0 was enabled as a third-party stand-alone annotator. All annotation outputs were manually inspected and curated. Gene maps were subsequently visualized as vector graphics using the OGDRAW^42^ web service (https://chlorobox.mpimp-golm.mpg.de/OGDraw.html).

Due to the large size and high repeat content of plant mitochondrial genomes, we used a de novo assembly strategy with PacBio CLR reads, followed by error correction using Illumina short reads. Initially, PacBio subreads were mapped to the complete mitochondrial genomes of closely related species using minimap2 (v2.28) with the -x map-pb option. As reference sequences, we used the mitochondrial genomes of *C. speciosa* (NCBI GenBank: AP029598.1), *C. avium* (NCBI RefSeq: NC_044768.1), *C.* × *kanzakura* (NC_065230.1), ‘Somei-yoshino’ (NC_065235.1), and *Prunus mume* (NC_065232.1). Subreads that mapped reliably to these references were extracted and assembled de novo using Canu, with parameters mostly set to default, except for genomeSize=400k, reflecting the expected mitochondrial genome size of *C. itosakura* and *C. jamasakura*. The resulting complete circular assemblies were polished using Illumina short reads. Mapping was performed with BWA-MEM (v0.7.18)^43^ under default settings, followed by error correction using Pilon (v1.24)^44^, also with default parameters. Gene annotation of the mitochondrial genomes was conducted using GeSeq, as with the chloroplast genome. While default configurations were largely maintained, specific adjustments were made: the input FASTA file was set as circular, the sequence source was designated as “mitochondrial,” and BLAT searches were performed using the mitochondrial genomes of ‘Somei-yoshino’, *C.* × *kanzakura*, *C. avium*, and *Arabidopsis thaliana* (NCBI RefSeq: NC_037304.1) as reference sequences. Annotation results were manually curated, and final gene maps were visualized as vector graphics using the OGDRAW web service, following the same procedure used for chloroplast genomes.

### Assembled genome evaluations

To evaluate the completeness and quality of the newly assembled genomes of *C. itosakura* and *C. jamasakura*, we employed multiple benchmarking approaches. BUSCO (v5.8.0)^45^ was used to assess the presence of conserved single-copy orthologs, with lineage datasets obtained from OrthoDB v10^46,47^. The most appropriate lineage dataset for each species was automatically selected using the --auto-lineage option, with “eudicots_odb10” tested for comparison. To complement and refine the BUSCO-based assessment, we employed Compleasm (v0.2.6)^48^, which offers a faster and more accurate estimation of genome completeness. Compleasm was run in run mode with the -l eudicots option.

Beyond ortholog-based evaluation, we assessed assembly quality using Inspector (v1.3.1)^49^, which evaluates structural completeness by aligning long reads back to the genome. In our analysis, PacBio CLR reads used in the assembly were aligned with the -d clr option to perform this assessment. Additionally, we applied a *k*-mer-based approach using Merqury (v1.3)^50^ to estimate consensus accuracy and completeness. *K*-mers were counted from PacBio CLR reads using Meryl (v1.3) with a *k*-mer size of 19. Merqury analyses were conducted under default settings to evaluate the pseudo-haploid assemblies, utilizing the Meryl-generated *k*-mer database.

### Assembled genome annotation

To annotate protein-coding genes, we first performed soft-masking of repetitive elements in the assembled genomes. For both *C. itosakura* and *C. jamasakura*, we conducted de novo identification of repetitive sequences using RepeatModeler2 with the -LTRStruct option enabled to incorporate LTR-specific detection pipelines. The resulting custom repeat libraries were then used with RepeatMasker to generate primary soft-masked assemblies, using the -xsmall option to mark repetitive regions in lowercase. Given that certain classes of tandem repeats are not fully captured by RepeatModeler2/RepeatMasker alone, we performed an additional round of soft-masking using Tandem Repeats Finder (TRF) (v4.09.1)^51^ to detect and mask these regions. TRF was executed with the following parameters: 2 7 7 80 10 50 500 -d -m -h. The tandem repeat regions identified by TRF were converted into BED format and merged with the previously masked assemblies using bedtools (v2.31.1)^52^, thereby producing a more comprehensively soft-masked genome suitable for gene prediction.

For structural annotation of protein-coding genes, we employed the “BRAKER with protein data” pipeline implemented in BRAKER3 (v3.0.8)^53^. As external protein evidence, we used the “Viridiplantae” partitioned protein dataset from OrthoDB v12^54^, downloaded from https://bioinf.uni-greifswald.de/bioinf/partitioned_odb12/. To enable both evidence-based and *ab initio* predictions, we activated the -- AUGUSTUS_ab_initio option, which allows AUGUSTUS to compute ab initio gene models in parallel with those supported by external protein hints. Furthermore, we specified the --busco_lineage option with “eudicots_odb10” to guide the parameter optimization of AUGUSTUS based on conserved gene structures within the eudicot lineage.

To infer the functional roles of genes predicted through BRAKER3-based structural annotation, we performed functional annotation using EnTAP (v2.2.0)^55^. EnTAP conducts homology-based searches using DIAMOND (v2.1.8)^56^, aligning predicted protein sequences against multiple reference databases, including SwissProt, TrEMBL, and the NCBI non-redundant (NR) plant protein database. Based on the results of EnTAP, we retained only those genes that were functionally annotated as belonging to Eukaryota, Viridiplantae, or Streptophyta.

To annotate repetitive elements, we utilized EDTA (v2.0.0)^57^, which integrates *de novo* and homology-based strategies to identify and classify transposable elements (TEs). For homology-based annotation, we incorporated the nrTEplants curated library (v0.3)^58^, a comprehensive collection of plant TEs compiled from multiple sources, including REdat^59^, RepetDB^60^, TREP^61^, and other specialized repositories. The EDTA analysis was conducted with the following parameters: --genome [genome.fasta] --cds [CDS.fasta] --curatedlib [nrTEplants curated library] --overwrite 1 --sensitive 1 --anno 1 --evaluate 1. To prevent misannotation of protein-coding regions as TEs, coding sequences (CDS) predicted by BRAKER3 were supplied as input. Each repetitive element was classified using RepeatMasker (v4.1.1), which is embedded in the EDTA Docker image (oushujun/edta:2.0.0).

Non-coding RNAs (ncRNAs) were annotated using a combination of specialized tools and databases targeting different ncRNA classes. Transfer RNAs (tRNAs) were predicted using tRNAscan-SE 2.0 (v2.0.12)^62^ with default parameters. Ribosomal RNAs (rRNAs) were identified using RNAmmer (v1.2)^63^ with the options -S euk -m lsu,ssu,tsu -multi, enabling the detection of large subunit (LSU), small subunit (SSU), and 5S rRNA genes in eukaryotic genomes. To annotate microRNAs (miRNAs) and other ncRNAs, we used Infernal (v1.1.4)^64^ with covariance models from Rfam 14.10^65^, including miRNA sequences curated in miRBase^66^. Infernal searches were conducted following Rfam’s recommended parameters, and an E-value threshold of “1e-10” was applied to retain only high-confidence hits.

As a final step, the genome assembly results, including structural features, GC content, gene density, distributions of non-coding RNAs, and annotated repetitive elements, were visualized using Circos (v0.69.9)^67^, a widely used circular genome visualization tool.

### Comparative genomics

For comparative genomic analysis, we collected reference genome assemblies and annotated protein sequences of *Prunus persica* (NCBI RefSeq: GCF_000346465.2), *C. campanulata* (NCBI GenBank: JBKKFS000000000.1), *C. speciosa* (NCBI GenBank: GCA_041154625.1), and two haplotypes of ‘Somei-yoshino’ (NCBI GenBank: GCA_005406145.1) from publicly available databases.

To investigate genome-wide synteny and structural variation, we constructed pairwise dot plots comparing each reference genome to the assemblies of *C. itosakura* and *C. jamasakura*. These comparisons were conducted using D-GENIES (v1.5.0)^68^, which performed sequence alignments internally with Minimap2 (v2.24) using the “Few repeats” setting to improve alignment accuracy in regions with low to moderate repeat content. We also performed interspecies synteny analysis using the protein sequences of *C. speciosa* and *C. campanulata*, which are among the most contiguous and complete assemblies available for closely related species.

To further characterize chromosomal-scale structural variation, we employed SyRI (v1.6.3)^69^. This tool analyzes whole-genome alignments to detect and classify a wide range of structural variants, including inversions, translocations, duplications, and indels. To compare the genomic distribution of non-coding RNAs, we used RIdeogram (v0.2.2)^70^, a visualization tool designed to generate idiograms overlaid with genomic features.

For genome-wide sequence similarity assessment, we used a combination of dnadiff from MUMmer4 and FastANI (v1.34)^71^. dnadiff, a utility within the MUMmer4 suite, utilizes nucmer alignments to compute detailed base-level statistics, including single nucleotide polymorphisms, insertions and deletions, and structural rearrangement breakpoints. To complement this alignment-based analysis, we also applied FastANI, which estimates average nucleotide identity through an alignment-free approach by comparing MinHash-based fragments.

To identify conserved syntenic blocks, we conducted all-versus-all homology searches of the predicted proteins using BLASTP (v2.12.0+)^27^ with an E-value cutoff of “1e–10”. The resulting alignment data were analyzed with MCScanX (v1.0.0)^72^, which detects collinear gene blocks across genomes. The synteny relationships identified by MCScanX were visualized using SynVisio (https://synvisio.github.io), enabling intuitive exploration of conserved gene order across species.

In addition, we conducted ortholog analysis using OrthoFinder (v2.5.5)^73,74^, which identifies orthogroups, reconstructs gene and species trees, and infers gene duplication events based on all-versus-all protein comparisons.

Reconstruction of Chromosome-Scale Haplotypes for ‘Somei-yoshino’ To enable direct comparison between the phased haplotypes of ‘Somei-yoshino’ and the wild-derived genomes of *C. speciosa* and *C. itosakura*, we reconstructed chromosome-level assemblies for each haplotype using RagTag. The phased contig assemblies of ‘Somei-yoshino’ were scaffolded separately: the *C. speciosa*-derived haplotype was scaffolded using our *C. speciosa* genome as the reference, while the *C. itosakura*-derived haplotype was scaffolded using our *C. itosakura* genome. This reference-guided scaffolding strategy allowed us to recover structurally resolved haplotypes of ‘Somei-yoshino’ that are directly comparable to their putative progenitor species, facilitating high-resolution structural and evolutionary analyses.

## Results

### Chromosome-level genome assemblies of *C. itosakura* and *C. jamasakura*

We assembled high-quality, chromosome-level genomes for *C. itosakura* and *C. jamasakura*, two phylogenetically and ecologically important wild cherry species. To provide a baseline for genome reconstruction, we first estimated genome size and heterozygosity using k-mer frequency distributions derived from Illumina short-read data. Haploid genome sizes were estimated to be approximately 229.81 Mbp for *C. itosakura* and 237.51 Mbp for *C. jamasakura*, with heterozygosity rates of 0.58% and 1.24%, respectively (Supplementary Fig. 2). These values reflect the genetic diversity inherent in wild, outcrossing populations and provided important parameters for guiding assembly and filtering.

The final nuclear genome assemblies were constructed through a multi-step approach involving long-read assembly, error correction, haplotig purging, organellar sequence removal, and reference-guided scaffolding. Details regarding the confidence of the reference-guided scaffolding are presented in Supplementary Fig. 3. For *C. itosakura*, the total assembled genome length was 259.08 Mbp, comprising 190 contigs with a contig N50 of 2.97 Mbp. Following chromosome-level scaffolding, the scaffold N50 increased to 30.63 Mbp. Similarly, the *C. jamasakura* assembly spanned 312.66 Mbp in total length, with 298 contigs and a contig N50 of 3.28 Mbp; the scaffold N50 after ordering and orienting was 32.39 Mbp. The overall GC content of the nuclear genome was calculated as 37.57% in *C. itosakura* and 37.84% in *C. jamasakura*. Each genome consisted of eight pseudomolecules corresponding to the haploid chromosome number (2n = 2x = 16), in line with cytological observations in *Cerasus* and other *Prunus* species (Fig. 2).

**Fig. 2.**
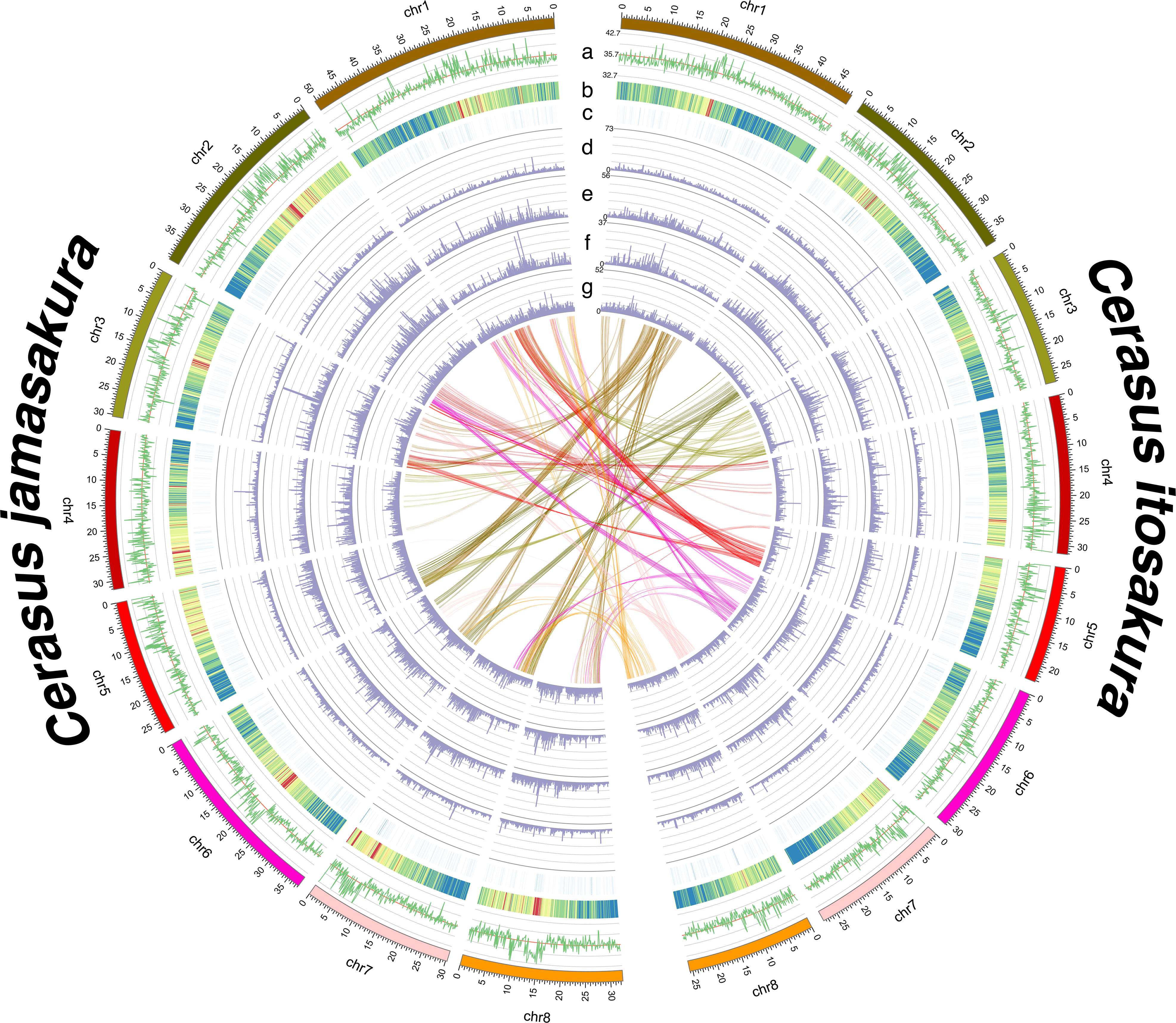
Schematic illustration of the genomic features of *C. itosakura* and *C. jamasakura*. The right half of the diagram corresponds to *C. itosakura* and the left half to *C. jamasakura*. From the outermost to the innermost track, each semicircle represents the following genomic characteristics: GC ratio (a), gene distribution (b), non-coding RNA distribution (c), long terminal repeats (LTR) (d), terminal inverted repeats (TIR) (e), non-LTR elements (LINE and SINE) (f), and non-TIR elements (Helitron) (g). Interchromosomal synteny between the *C. itosakura* and *C. jamasakura* genomes is shown at the center of the figure.

To rigorously assess the completeness, accuracy, and structural integrity of the assembled genomes, we conducted multiple complementary benchmarking analyses. Using the eudicot dataset from BUSCO, we identified 98.75% completeness with 90.84% (n=2113) complete single-copy orthologs, 7.91% (n=184) duplicated, 0.64% (n=15) fragmented, and 0.60% (n=14) missing in *C. itosakura*. Similarly, the *C. jamasakura* assembly contained 98.62% completeness with 79.41% (n=1847) single-copy, 19.22% (n=447) duplicated, 0.64% (n=15) fragmented, and 0.73% (n=17) missing orthologs. These results indicate near-complete representation of the conserved gene space in both species, with minimal gene loss or fragmentation. Compleasm analysis, which provides a fast and highly concordant alternative to BUSCO, corroborated these findings. The *C. itosakura* genome showed 99.48% completeness with 92.00% (n=2140) single-copy, 7.48% (n=174) duplicated, 0.04% (n=1) fragmented, and 0.47% (n=11) missing genes, while the *C. jamasakura* assembly yielded 99.48% completeness with 80.65% (n=1876) single-copy, 18.83% (n=438) duplicated, 0.04% (n=1) fragmented, and 0.47% (n=11) missing, further supporting the reliability and completeness of the assemblies. To evaluate consensus-level sequence accuracy, we employed Merqury analysis, which estimates genome quality based on *k*-mer concordance between the reads and the assembly. The quality value (QV) scores were 44.86 for *C. itosakura* and 42.35 for *C. jamasakura*, with estimated completeness values of 93.62% and 88.25%, respectively. These metrics suggest that the majority of true genomic content was accurately reconstructed and retained. In addition, long-read alignment-based evaluation using Inspector revealed high structural integrity for both genomes. The read mapping rates exceeded 88.08% for *C. itosakura* and 91.46% for *C. jamasakura*, with QV scores 31.93 and 30.66, respectively. Together, these evaluations confirm that both assemblies are of high quality in terms of gene content, base-level accuracy, and structural continuity, making them suitable for downstream analyses such as gene annotation, comparative genomics, and evolutionary inference.

### Organelle genome structures and gene content

We successfully assembled and annotated complete organelle genomes for *C. itosakura* and *C. jamasakura*, including both chloroplast and mitochondrial DNA. The chloroplast genomes of both species exhibited the typical quadripartite structure found in most angiosperms, composed of a large single-copy (LSC) region, a small single-copy (SSC) region, and two inverted repeats (IRs). The total lengths of the plastomes were 157.81 Kbp in *C. itosakura* and 157.90 Kbp in *C. jamasakura*. Gene annotation identified 88 protein-coding genes, 37 tRNAs, and 8 rRNAs, with no evidence of gene loss or structural rearrangement compared to other *Cerasus* species (Fig. 3). The overall gene order and content were highly conserved, suggesting a stable chloroplast architecture across the lineage.

**Fig. 3.**
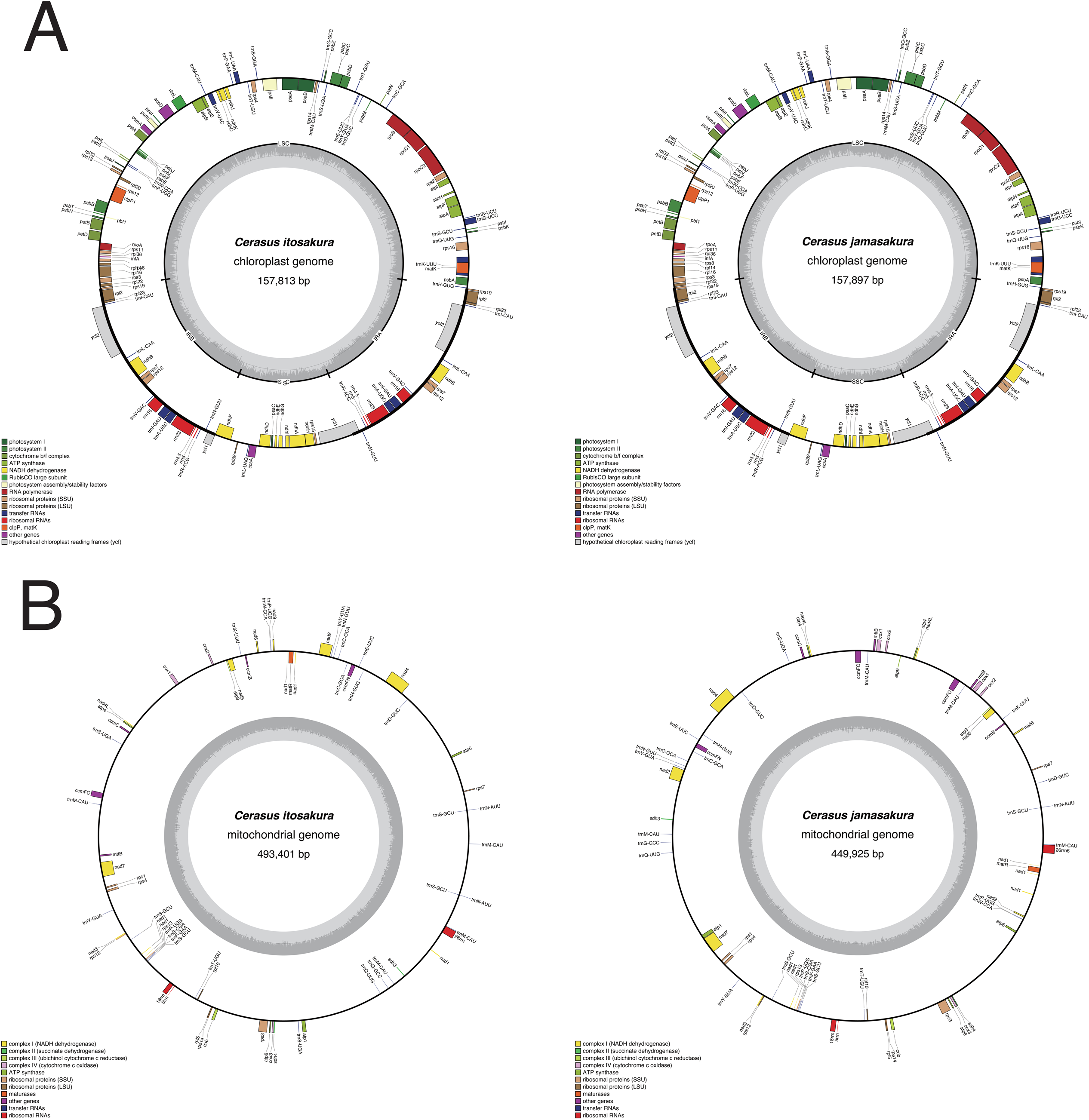
Schematic diagrams of gene annotations for the chloroplast and mitochondrial genomes of *C. itosakura* and *C. jamasakura*. Panel A shows the chloroplast genomes, with *C. itosakura* on the left and *C. jamasakura* on the right. Panel B shows the mitochondrial genomes, with *C. itosakura* on the left and *C. jamasakura* on the right. In each diagram, the positional relationships among genes within the circular genomes are illustrated, and gene types are indicated in the corresponding legends. The innermost track represents the GC ratio (darker color). For the chloroplast genomes, the small single-copy (SSC), large single-copy (LSC), and inverted repeat regions (IRA and IRB) are depicted in the inner circle.

The mitochondrial genomes of both species were also assembled into complete circular molecules, with total lengths of 493.40 Kbp in *C. itosakura* and 449.93 Kbp in *C. jamasakura*. As is typical for plant mitochondria, these assemblies displayed substantially greater structural complexity and a higher repeat content than their plastid counterparts. We identified 35 and 42 protein-coding genes in *C*. *itosakura* and *C. jamasakura*, respectively. These included essential components of the electron transport chain and ribosomal machinery. Additionally, we detected 29 and 27 tRNAs, and 3 and 3 rRNAs in the respective species. Several lineage-specific differences were observed, including repeat-mediated expansions and intron variants, reflecting the dynamic and recombinogenic nature of plant mitochondrial genomes (Fig. 3).

### Annotation of coding and non-coding elements

To characterize the gene content and regulatory landscape of the assembled genomes, we performed comprehensive annotations of both coding and non-coding elements in *C. itosakura* and *C. jamasakura*. We first identified and masked repetitive sequences using a combination of de novo and homology-based approaches. The total repeat content accounted for 43.10% of the *C. itosakura* genome and 46.91% of the *C. jamasakura* genome, with long terminal repeat (LTR) retrotransposons being the predominant class in both species. Ty3/Gypsy elements were particularly abundant, followed by Ty1/Copia elements, consistent with observations in other Prunus genomes. Notably, the relative proportion of LTRs differed modestly between the two species, suggesting lineage-specific variation in recent transpositional activity.

We then conducted structural annotation of protein-coding genes. The initial gene prediction yielded 39,062 gene models in *C. itosakura* and 37,751 in *C. jamasakura* based on evidence-guided *ab initio* modeling. To refine these predictions and reduce false positives, we applied a filtering step in which only gene models supported by external protein homology evidence were retained. As a result, 34,516 and 33,570 protein-coding genes were ultimately retained as high-confidence annotations in *C. itosakura* and *C. jamasakura*, respectively. These filtered gene sets were used for downstream structural and functional analyses. The annotated genes displayed structural characteristics consistent with those of other eudicots. The average number of exons per gene was 5.19 in *C. itosakura* and 5.42 in *C. jamasakura*, while the average coding sequence (CDS) lengths were 1248.54 bp and 1272.03 bp, respectively.

We also annotated a broad array of non-coding RNAs (ncRNAs) in each genome. In *C. itosakura*, we identified 608 transfer RNAs (tRNAs), 234 ribosomal RNAs (rRNAs), and 279 predicted microRNAs (miRNAs), along with additional small RNAs corresponding to snoRNAs and other Rfam families. In *C. jamasakura*, the genome encoded 677 tRNAs, 318 rRNAs, and 346 putative miRNAs, along with similar classes of small RNAs. While the overall ncRNA content was broadly conserved, certain differences in copy number and chromosomal localization were observed between the two species, particularly in the distribution of rRNAs. Several rRNA loci appeared to be lineage-specific, suggesting lineage-specific rearrangements or differential retention/loss events during genome evolution (Fig. 4).

**Fig. 4.**
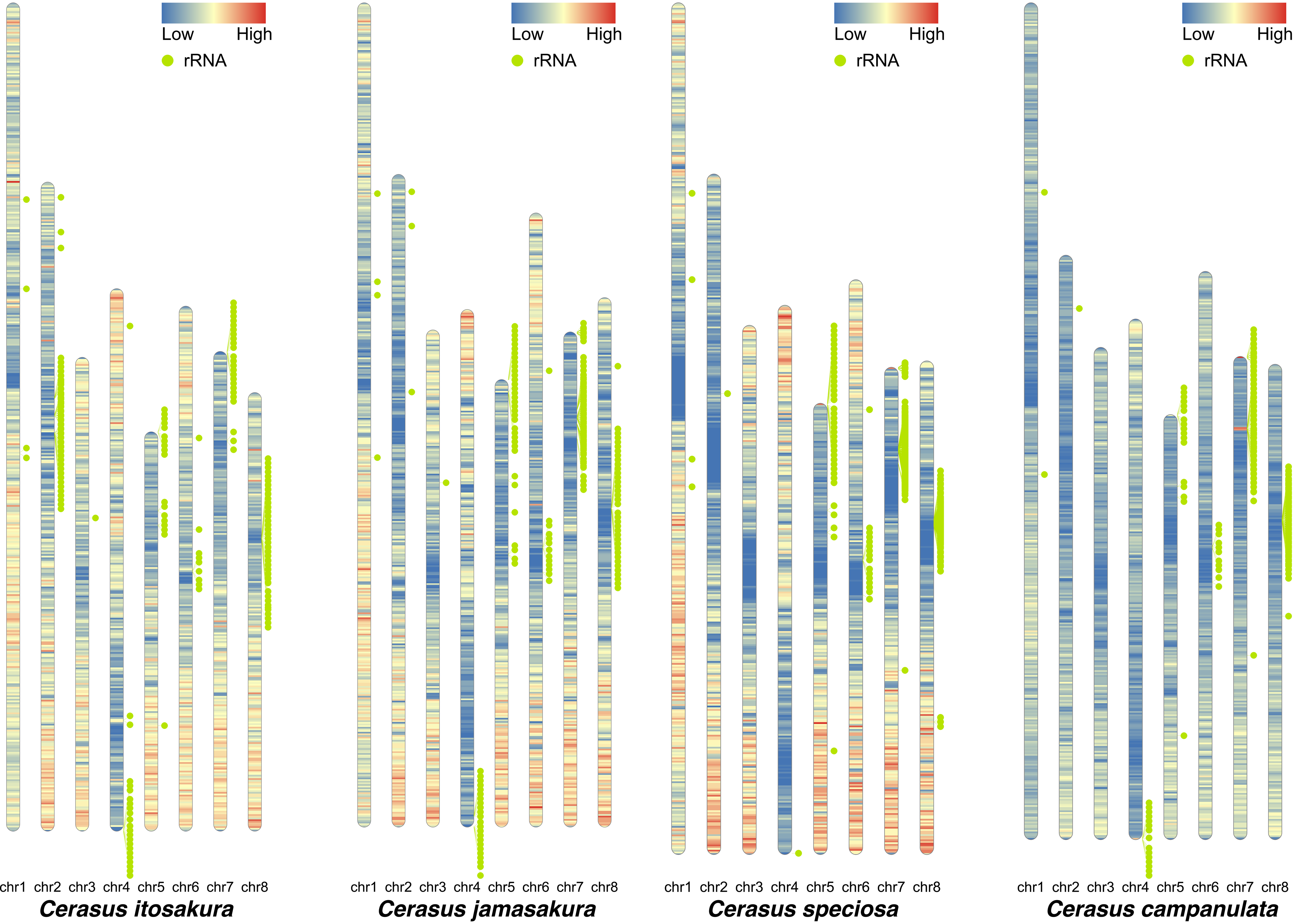
Species-specific differences were observed in the chromosomal distribution of rRNA gene clusters. In addition to the two newly assembled genomes, the genomes of *C. speciosa* and *C. campanulata* were included for comparison. Each bar represents a karyotype, with the color gradient indicating gene density: red corresponds to higher density, whereas blue corresponds to lower density. Green circles represent rRNA genes (rDNA), and the green lines extending from them indicate their positions on the karyotype. While some rDNA clusters are located in positions shared across all species, others appear to be species-specific.

### Comparative genomics and genome evolution in *Cerasus*

To quantify the overall sequence similarity among *Cerasus* species, we performed both whole-genome alignment and average nucleotide identity (ANI) analyses. These analyses included *C. itosakura*, *C. jamasakura*, *C. speciosa*, *C. campanulata*, and *Prunus persica* as an outgroup. Whole-genome alignments were conducted using *dnadiff* implemented in Mummer4, yielding one-to-one and many-to-many alignment statistics for each species pair. These results provided detailed measurements of aligned sequence lengths, percent identity, SNP counts, indels, and large-scale structural breakpoints. The highest sequence-level similarity was observed between *C. jamasakura* and *C. speciosa*, indicating a particularly close genomic relationship (Table 1). Complementary ANI analyses further supported this pattern: the highest ANI was also found between *C. jamasakura* and *C. speciosa*, with *C. itosakura* exhibiting slightly lower but still high ANI values with both species. Complete alignment and ANI results are summarized in Supplementary Table 1.

**Table 1.**
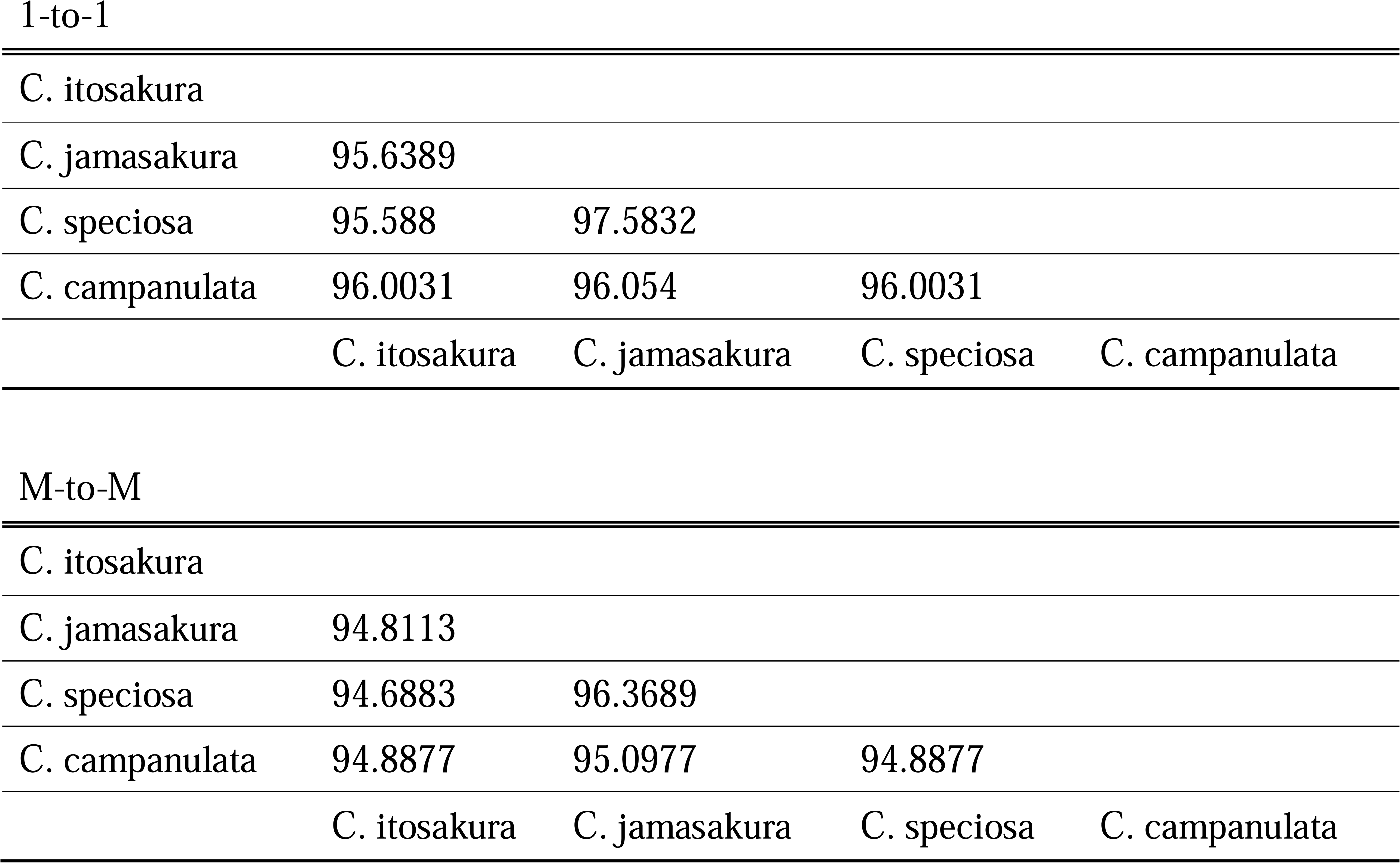
Genome difference in genus Cerasus.

1-to-1
|  |  |  |  |  |
| --- | --- | --- | --- | --- |
| C. itosakura |  |  |  |  |
| C. jamasakura | 95.6389 |  |  |  |
| C. speciosa | 95.588 | 97.5832 |  |  |
| C. campanulata | 96.0031 | 96.054 | 96.0031 |  |
|  | C. itosakura | C. jamasakura | C. speciosa | C. campanulata |
| M-to-M |  |  |  |  |
| C. itosakura |  |  |  |  |
| C. jamasakura | 94.8113 |  |  |  |
| C. speciosa | 94.6883 | 96.3689 |  |  |
| C. campanulata | 94.8877 | 95.0977 | 94.8877 |  |
|  | C. itosakura | C. jamasakura | C. speciosa | C. campanulata |

Despite the overall genomic similarity, dot plot analyses revealed several large-scale structural rearrangements among *C. itosakura*, *C. jamasakura*, and *C. campanulata*, using *C. speciosa* as the reference genome (Supplementary Fig. 4). While genome-wide chromosomal collinearity was generally conserved, multiple inversions were detected, as summarized in a histogram of inversion lengths (Supplementary Fig. 5). Among these, two species-specific inversions exceeding 1 Mb in length were notable: one on chromosome 2 of *C. speciosa* and the other on chromosome 8 of *C. itosakura* (Fig. 5). The former likely represents a lineage-specific rearrangement in *C. speciosa*, and was not further investigated due to its limited relevance to the focal species of this study.

**Fig. 5.**
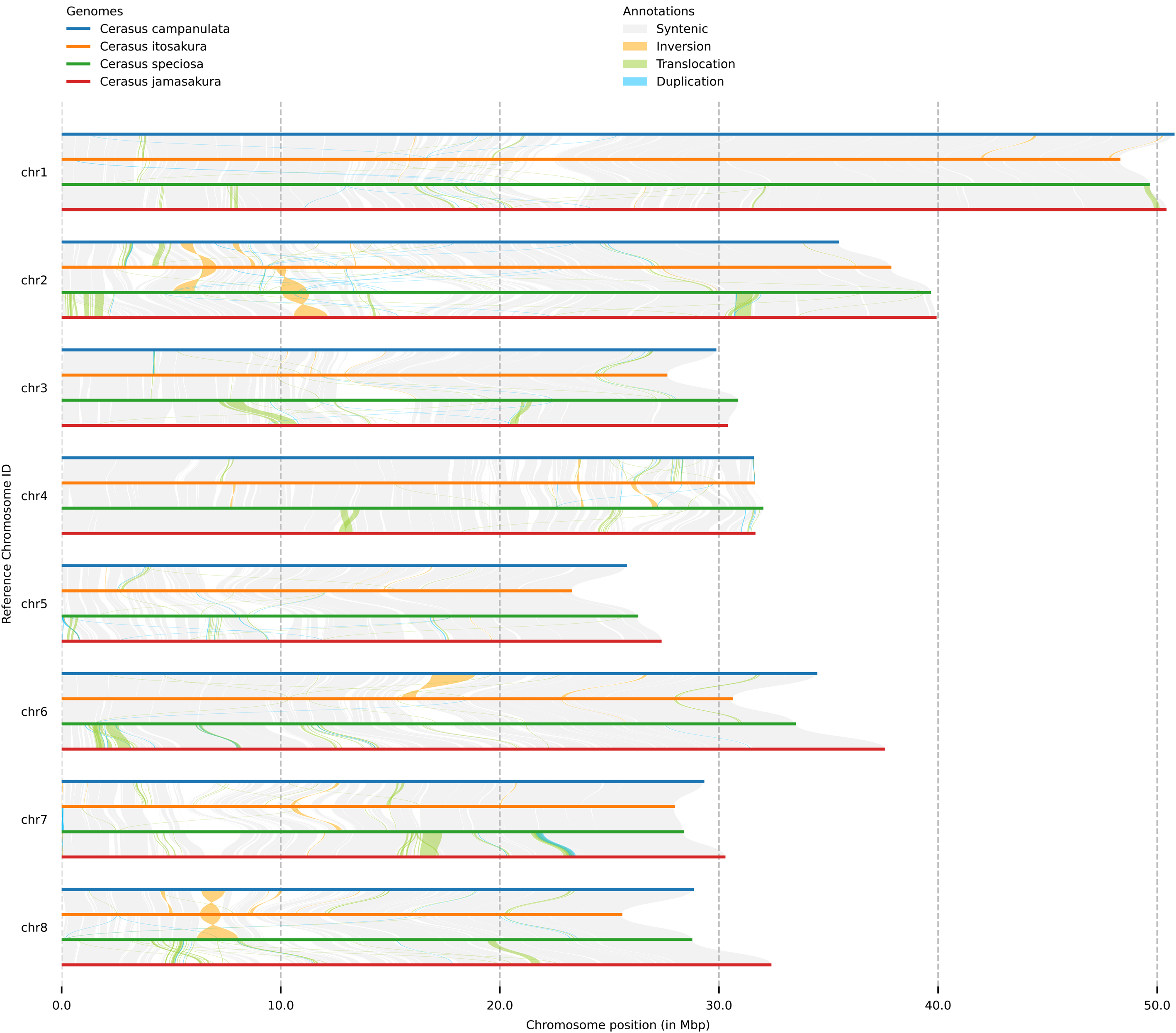
Chromosome-scale synteny among four *Cerasus* species. Comparative chromosome alignments of *C. campanulata*, *C. itosakura*, *C. speciosa*, and *C. jamasakura*. Alignments are color-coded according to structural relationships, including conserved synteny, inversions, translocations, and duplications.

In contrast, the inversion on chromosome 8 of *C. itosakura* was absent in both *C. jamasakura* and other *Cerasus* genomes, indicating its specificity to *C. itosakura*. Importantly, the rearranged region was contained within a single contig, excluding the possibility of misassembly during reference-guided scaffolding. This inversion spans 1,840,090 bp and includes 82 protein-coding genes, corresponding to 88 mRNA isoforms, which are listed in Supplementary Table 2.

We also examined gene collinearity across the four *Cerasus* genomes, *C. itosakura*, *C. jamasakura*, *C. speciosa*, and *C. campanulata*, to evaluate the conservation of gene order. The analysis revealed extensive syntenic relationships, reflecting the close evolutionary proximity among these species.

Comparison of orthologous gene sets revealed that 67.0% of genes were shared across the four Cerasus genomes (*C. itosakura*, *C. jamasakura*, *C. speciosa*, and *C. campanulata*), indicating a substantial core gene repertoire within the genus. When the comparison was extended to include *P. persica* as an outgroup, the proportion of shared genes decreased to 63.1%. Both results are presented in Supplementary Fig. 6.

Chromosome-scale comparison of ‘Somei-yoshino’ haplotypes with wild progenitors To investigate the genomic composition and structural integrity of the interspecific hybrid ‘Somei-yoshino,’ we reconstructed its haplotype-resolved, chromosome-scale genome by applying reference-guided scaffolding to the publicly available contig-level assembly. This was performed separately for each haplotype using our newly generated high-quality genomes of *C. itosakura* and *C. speciosa*, the two wild progenitor species hypothesized to constitute the parental origins of ‘Somei-yoshino.’ Details regarding the confidence of the reference-guided scaffolding for ‘Somei-yoshino’ are presented in Supplementary Fig. 7.

Dot plot analyses between the scaffolded ‘Somei-yoshino’ haplotypes and their respective wild progenitor genomes revealed extensive collinearity and no apparent large-scale structural rearrangements (Supplementary Fig. 8). Complementary whole-chromosome alignments further supported this conclusion, showing no major structural variations (Supplementary Fig. 9). Notably, however, the total genome length of the public ‘Somei-yoshino’ assembly was substantially shorter than those of the wild progenitors, suggesting that portions of genomic content may be missing from the existing assembly.

To assess the authenticity of the *C. speciosa*-derived haplotype in ‘Somei-yoshino’, we compared the grouping confidence of contigs scaffolded using either the *C. speciosa* or *C. jamasakura* genome. Mann-Whitney U tests indicated significantly higher grouping confidence (*p-*value: 0.011775) when using *C. speciosa* as the reference (mean = 0.959140, SE = 0.005272), compared to when using *C. jamasakura* (mean = 0.937931, SE = 0.006630), lending quantitative support to the hypothesis that this haplotype originated from *C. speciosa*. This distinction is particularly relevant given the close phylogenetic relationship between *C. speciosa* and *C. jamasakura*.

Finally, we evaluated sequence similarity between each ‘Somei-yoshino’ haplotype and the corresponding wild progenitor genomes using dnadiff from the MUMmer4 package and average nucleotide identity (ANI) metrics. Neither haplotype showed perfect identity with its putative progenitor, and the *C. speciosa* haplotype in particular exhibited lower sequence similarity than expected (Supplementary Table 3 and 4). These findings suggest that while ‘Somei-yoshino’ retains broad genomic similarity to its progenitors, both haplotypes may have undergone post-hybridization sequence divergence or assembly-related data loss. In addition to these possibilities, intraspecific polymorphism should also be considered as a potential source of the observed discrepancies.

## Discussion

We report the first chromosome-scale genome assemblies of *C. itosakura* and *C. jamasakura*, two ecologically and phylogenetically important wild cherry species that previously lacked high-quality reference genomes. These assemblies close a significant gap in the genomic resources for *Cerasus* and establish a solid foundation for comparative and functional genomic research across cherry and related *Prunus* species. Both genomes demonstrate exceptional completeness (BUSCO ≥98.6%) and high base-level accuracy (Merqury QV >42), ensuring that the conserved gene space is nearly fully represented with minimal errors. With scaffold N50 values approaching 30 Mb, the assemblies exhibit remarkable contiguity, rivaling the best available reference genomes for wild cherries. Importantly, all eight pseudomolecules corresponding to the haploid chromosome complement (2n = 2x = 16) were successfully reconstructed^36–39^, underscoring the reliability of our sequencing and scaffolding strategy.

Our comparative genomic analysis of four *Cerasus* species revealed pronounced, species-specific differences in the chromosomal distribution of rRNA gene clusters. Such divergence, even among closely related taxa, has been reported in other groups and is thought to reflect rapid evolutionary change driven by transposition and other rearrangement processes ^75,76^. Intragenomic variation in rDNA copy number and chromosomal localization can arise from unequal crossing-over and may be further shaped by the activity of transposable elements^76,77^. In plants, rDNA sites frequently occupy non-random positions such as terminal or pericentromeric regions, and shifts in their distribution are often associated with lineage-specific chromosomal restructuring events^78,79^. The distinct patterns observed here suggest that similar mechanisms have contributed to the diversification of rDNA organization within *Cerasus* lineages, making rDNA loci valuable markers for tracing chromosomal and evolutionary dynamics.

One of the most striking findings of this study is the identification of a large, species-specific chromosomal inversion in *C. itosakura*. This ∼1.84 Mb inversion on chromosome 8 is absent from the genomes of *C. jamasakura*, *C. speciosa*, *C. campanulata*, and other analyzed cherry species. The rearranged segment, fully resolved within a single contig, contains 82 protein-coding genes (88 transcripts), indicating that the inversion is a bona fide structural variant rather than a misassembly. Such large inversions can have important evolutionary consequences by suppressing recombination in heterozygotes, facilitating the maintenance of linked gene complexes, and potentially contributing to reproductive isolation or local adaptation^80–83^. Although the phenotypic impact of this inversion remains to be elucidated, its presence suggests a previously unrecognized genomic divergence unique to the *C. itosakura* lineage. Future population-scale investigations will be crucial to determine its prevalence and adaptive relevance.

Beyond the identified inversion, our analyses revealed strong genome-wide collinearity and high sequence identity across the wild cherry genomes. Whole-genome alignment and average nucleotide identity (ANI) analyses demonstrated that *C. jamasakura* is most closely related to *C. speciosa*, consistent with their taxonomic classification within the core group of Japanese flowering cherries. This group represents the genetic foundation of many ornamental cultivars that have arisen through natural and artificial hybridization among approximately ten *Cerasus* species in Japan. The availability of a high-quality genome assembly for *C. jamasakura* now provides a definitive reference for one of these key taxa, enabling precise comparative analyses and improved resolution of species boundaries.

In addition to nuclear genome assemblies, we generated mitochondrial genome sequences based on long-read data. These assemblies reflect one circular configuration inferred from the data; however, plant mitochondrial genomes are highly dynamic and structurally complex in vivo, typically existing as a mixture of subgenomic molecules and recombination-driven isoforms^84–89^. Thus, our assemblies should be interpreted as consensus representations of one possible structural state rather than exhaustive reconstructions of the in vivo mitochondrial architecture. Nonetheless, they provide a useful basis for comparative and functional mitochondrial analyses.

Beyond the large inversion in *C. itosakura*, we found no evidence of major structural variation among the analyzed wild genomes. This suggests that karyotypic structure has remained largely conserved during the diversification of flowering cherries, consistent with their shared chromosome number (2n = 2x = 16). Taken together, our results support the taxonomic distinction between *C. itosakura* and *C. jamasakura* as separate species while reinforcing their shared placement within the broader *Prunus/Cerasus* clade. These new genomic resources open avenues for addressing fundamental questions in cherry evolution, diversification, and domestication, and provide a critical toolkit for future work in conservation biology, evolutionary genomics, and breeding.

The new reference genomes also facilitated refined genomic characterization of the interspecific hybrid ‘Somei-yoshino’, Japan’s iconic ornamental cherry. By anchoring its haplotype-resolved genome to the *C. itosakura* and *C. speciosa* references, we confirmed its hybrid origin at the chromosome level. Each haplotype exhibited strong collinearity with one of the parental genomes, with no evidence of major structural rearrangements. Notably, the *C. speciosa*-derived haplotype aligned more closely with our *C. speciosa* reference than with *C. jamasakura*, providing clear molecular support for the longstanding hypothesis that *C. speciosa* served as the paternal progenitor^5,18^. Minor reductions in sequence identity between the hybrid haplotypes and their wild ancestors may reflect post-hybridization divergence or assembly artifacts, but the overall structural conservation supports previous reports of genomic stability among closely related cherries. In addition, intraspecific polymorphism cannot be excluded as a potential contributor to these differences. These results underscore the importance of high-quality references for resolving hybrid ancestry and genome composition in domesticated or ornamental taxa.

### Conclusion

In summary, this work represents a major advance in cherry genomics. We generated the first chromosome-level reference genomes for *C. itosakura* and *C. jamasakura*, significantly expanding the genomic resources available for wild Cerasus species. These assemblies are highly complete and structurally accurate, capturing nearly the full complement of genes and faithfully reconstructing chromosomal architecture. Among the key findings is a large, lineage-specific inversion in *C. itosakura*, providing new insight into structural divergence within the genus. Comparative analyses clarified evolutionary relationships and revealed strong genome-wide collinearity among wild cherries, while also confirming the hybrid origin of ‘Somei-yoshino’ at the chromosome level. Collectively, these genomic resources and discoveries deepen our understanding of cherry genome evolution and species diversification, and offer a robust foundation for downstream applications in taxonomy, horticulture, breeding, and conservation.

## Acknowledgments

We thank Motoko Nihei and Kumiko Masuyama for their technical assistance in this study.

## Data availability

The sequencing datasets generated and utilized in this study have been deposited in public repositories to ensure open access and reproducibility. Long-read and short-read sequencing data for both *C. itosakura* and *C. jamasakura* are available from the DNA Data Bank of Japan (DDBJ) Sequence Read Archive (DRA). Specifically, PacBio reads are accessible under accession numbers DRR689950 (*C. itosakura*) and DRR689952 (*C. jamasakura*), while Illumina reads are registered as DRR689951 and DRR689953 for the respective species. The genome assemblies have been submitted to the DDBJ under the accession identifiers AP041240–AP041289 for *C. itosakura* and AP041290– AP041398 for *C. jamasakura*. Functional annotations, including GFF files and other genome feature data, have been made publicly available through Figshare at 10.6084/m9.figshare.30000994.

## Reference

1. Arakawa, H. 1955, Twelve centuries of blooming dates of the cherry blossoms at the city of Kyoto and its own vicinity. Geofisica Pura e Applicata, 30, 147–150.

2. Kuitert, W. and Peterse, A. 1999, Japanese flowering cherries. Timber Press: Oregon.

3. Aono, Y. and Kazui, K. 2007, Phenological data series of cherry tree flowering in Kyoto, Japan, and its application to reconstruction of springtime temperatures since the 9th century. International Journal of Climatology, 28, 905–914.

4. Jefferson, R. M. and Wain, K. K. 1984, The nomenclature of cultivated Japanese flowing cherries (Prunus): the Sato-zakura group. U.S. Dept. of Agriculture, Agricultural Research Service: [Washington, D.C.].

5. Kato, S., Matsumoto, A., Yoshimura, K., et al. 2014, Origins of Japanese flowering cherry (Prunus subgenus Cerasus) cultivars revealed using nuclear SSR markers. Tree Genetics & Genomes, 10, 477–487.

6. Ohba, H. 2001, Cerasus. In: Iwatsuki, K., Yamazaki, T., Boufford, D. E. and Ohba, H. (eds), Flora of Japan, Kodansha, Tokyo, pp. 128–144.

7. Kawasaki, T. 1993, Flowering cherries of Japan. Yama-Kei Publisher: Tokyo.

8. Ohba, H., Kawasaki, T., Kihara, H. and Tanaka, H. 2007, Flowering cherries of Japan. Yama-Kei Publishers: Tokyo.

9. Ikeda, H., Iketani, H. and Katsuki, T. 2017, Rosaceae. In: Ohashi, H., Kadota, Y., Murata, J., Yonekura, K. and Kihara, H. (eds), Wild Flowers of Japan, revised new edition, Heibonsha, Tokyo, pp. 23–88.

10. Katsuki, T. 2018, A New Species, Cerasus kumanoensis from the Southern Kii Peninsula, Japan. Acta Phytotaxonomica et Geobotanica, 69, 119–126.

11. Katsuki, T. and Iketani, H. 2016, Nomenclature of Tokyo cherry (Cerasus × yedoensis ‘Somei - yoshino’, Rosaceae) and allied interspecific hybrids based on recent advances in population genetics. Taxon, 65, 1415–1419.

12. Innan, H., Terauchi, R., Miyashita, N. T. and Tsunewaki, K. 1995, DNA fingerprinting study on the intraspecific variation and the origin of Prunus yedoensis (Someiyoshino). The Japanese Journal of Genetics, 70, 185–196.

13. Takenaka, Y. 1963, The origin of the yoshino cherry tree. The Journal of Heredity, 54, 207–211.

14. Iketani, H., Ohta, S., Kawahara, T., et al. 2007, Analyses of clonal status in ‘Somei-yoshino’ and confirmation of genealogical record in other cultivars of Prunus ×yedoensis by microsatellite markers. Breeding Science, 57, 1–6.

15. Nakamura, I., Takahashi, H., Ohta, S., et al. 2015, Origin of Prunus x yedoenins ‘Somei-yoshino’ based on sequence analysis of PolA1 gene. Advances in Horticultural Science, 29, 17–23.

16. Fujiwara, K., Toyoda, A., Biswa, B. B., et al. 2025, A near complete genome assembly of the Oshima cherry Cerasus speciosa. Scientific Data, 12, 162.

17. Jiang, D., Li, Y., Zhuge, F., et al. 2025, The telomere-to-telomere genome of flowering cherry (Prunus campanulata) reveals genomic evolution of the subgenus Cerasus. Gigascience, 14.

18. Shirasawa, K., Esumi, T., Hirakawa, H., et al. 2019, Phased genome sequence of an interspecific hybrid flowering cherry, ‘Somei-Yoshino’ (Cerasus × yedoensis). DNA Res, 26, 379–389.

19. Shi, S., Li, J., Sun, J., Yu, J. and Zhou, S. 2013, Phylogeny and classification of Prunus sensu lato (Rosaceae). Journal of Integrative Plant Biology, 55, 1069–1079.

20. Zhang, J., Wang, Y., Chen, T., et al. 2021, Evolution of Rosaceae plastomes highlights unique Cerasus diversification and independent origins of fruiting cherry. Frontiers in Plant Science, 12, 736053.

21. Wan, T., Qiao, B.-x., Zhou, J., et al. 2023, Evolutionary and phylogenetic analyses of 11 Cerasus species based on the complete chloroplast genome. Frontiers in Plant Science, 14, 1070600.

22. Chen, S., Zhou, Y., Chen, Y. and Gu, J. 2018, fastp: an ultra-fast all-in-one FASTQ preprocessor. Bioinformatics, 34, i884–i890.

23. Kokot, M., Dlugosz, M. and Deorowicz, S. 2017, KMC 3: counting and manipulating k-mer statistics. Bioinformatics, 33, 2759–2761.

24. Ranallo-Benavidez, T. R., Jaron, K. S. and Schatz, M. C. 2020, GenomeScope 2.0 and Smudgeplot for reference-free profiling of polyploid genomes. Nat Commun, 11, 1432.

25. Vurture, G. W., Sedlazeck, F. J., Nattestad, M., et al. 2017, GenomeScope: fast reference-free genome profiling from short reads. Bioinformatics, 33, 2202–2204.

26. Koren, S., Walenz, B. P., Berlin, K., Miller, J. R., Bergman, N. H. and Phillippy, A. M. 2017, Canu: scalable and accurate long-read assembly via adaptive k-mer weighting and repeat separation. Genome Res, 27, 722–736.

27. Altschul, S. F., Gish, W., Miller, W., Myers, E. W. and Lipman, D. J. 1990, Basic local alignment search tool. Journal of Molecular Biology, 215, 403–410.

28. Laetsch, D. R. and Blaxter, M. L. 2017, BlobTools: Interrogation of genome assemblies. F1000Research, 6.

29. Li, H. 2018, Minimap2: pairwise alignment for nucleotide sequences. Bioinformatics, 34, 3094–3100.

30. Roach, M. J., Schmidt, S. A. and Borneman, A. R. 2018, Purge Haplotigs: allelic contig reassignment for third-gen diploid genome assemblies. BMC Bioinformatics, 19, 460.

31. Flynn, J. M., Hubley, R., Goubert, C., et al. 2020, RepeatModeler2 for automated genomic discovery of transposable element families. Proc Natl Acad Sci U S A, 117, 9451–9457.

32. Smit, A. F. A., Hubley, R. and Green, P. RepeatMasker.

33. Marcais, G., Delcher, A. L., Phillippy, A. M., Coston, R., Salzberg, S. L. and Zimin, A. 2018, MUMmer4: A fast and versatile genome alignment system. PLoS Comput Biol, 14, e1005944.

34. Hu, J., Fan, J., Sun, Z. and Liu, S. 2020, NextPolish: a fast and efficient genome polishing tool for long-read assembly. Bioinformatics, 36, 2253–2255.

35. Alonge, M., Lebeigle, L., Kirsche, M., et al. 2022, Automated assembly scaffolding using RagTag elevates a new tomato system for high-throughput genome editing. Genome Biol, 23, 258.

36. Oginuma, K. and Tanaka, R. 1976, Karyomorphological studies on some cherry trees in Japan. The Journal of Japanese Botany, 51, 104–109.

37. Iwatsubo, Y., Kawasaki, T. and Naruhashi, N. 2003, Chromosome numbers of 41 cultivated taxa of Prunus subg. Cerasus in Japan. Journal of Phytogeography and Taxonomy, 51, 165–168.

38. Iwatsubo, Y., Kawasaki, T. and Naruhashi, N. 2002, Chromosome numbers of 193 cultivated taxa of Prunus subg. Cerasus in Japan. Journal of Phytogeography and Taxonomy, 50, 21–34.

39. Iwatsubo, Y., Sengi, Y. and Naruhashi, N. 2004, Chromosome numbers of 36 cultivated taxa of Prunus subg. Cerasus in Japan. Journal of Phytogeography and Taxonomy, 52, 73–76.

40. Jin, J. J., Yu, W. B., Yang, J. B., et al. 2020, GetOrganelle: a fast and versatile toolkit for accurate de novo assembly of organelle genomes. Genome Biol, 21, 241.

41. Tillich, M., Lehwark, P., Pellizzer, T., et al. 2017, GeSeq - versatile and accurate annotation of organelle genomes. Nucleic Acids Res, 45, W6–W11.

42. Greiner, S., Lehwark, P. and Bock, R. 2019, OrganellarGenomeDRAW (OGDRAW) version 1.3.1: expanded toolkit for the graphical visualization of organellar genomes. Nucleic Acids Res, 47, W59–W64.

43. Li, H. and Durbin, R. 2009, Fast and accurate short read alignment with Burrows-Wheeler transform. Bioinformatics, 25, 1754–1760.

44. Walker, B. J., Abeel, T., Shea, T., et al. 2014, Pilon: an integrated tool for comprehensive microbial variant detection and genome assembly improvement. PLoS One, 9, e112963.

45. Manni, M., Berkeley, M. R., Seppey, M., Simao, F. A. and Zdobnov, E. M. 2021, BUSCO Update: Novel and Streamlined Workflows along with Broader and Deeper Phylogenetic Coverage for Scoring of Eukaryotic, Prokaryotic, and Viral Genomes. Mol Biol Evol, 38, 4647–4654.

46. Kriventseva, E. V., Kuznetsov, D., Tegenfeldt, F., et al. 2019, OrthoDB v10: sampling the diversity of animal, plant, fungal, protist, bacterial and viral genomes for evolutionary and functional annotations of orthologs. Nucleic Acids Res, 47, D807–D811.

47. Zdobnov, E. M., Kuznetsov, D., Tegenfeldt, F., Manni, M., Berkeley, M. and Kriventseva, E. V. 2021, OrthoDB in 2020: evolutionary and functional annotations of orthologs. Nucleic Acids Res, 49, D389–D393.

48. Huang, N. and Li, H. 2023, compleasm: a faster and more accurate reimplementation of BUSCO. Bioinformatics, 39.

49. Chen, Y., Zhang, Y., Wang, A. Y., Gao, M. and Chong, Z. 2021, Accurate long-read de novo assembly evaluation with Inspector. Genome Biol, 22, 312.

50. Rhie, A., Walenz, B. P., Koren, S. and Phillippy, A. M. 2020, Merqury: reference-free quality, completeness, and phasing assessment for genome assemblies. Genome Biology, 21.

51. Benson, G. 1999, Tandem repeats finder: a program to analyze DNA sequences. Nucleic Acids Res, 27, 573–580.

52. Quinlan, A. R. and Hall, I. M. 2010, BEDTools: a flexible suite of utilities for comparing genomic features. Bioinformatics, 26, 841–842.

53. Gabriel, L., Bruna, T., Hoff, K. J., et al. 2024, BRAKER3: Fully automated genome annotation using RNA-seq and protein evidence with GeneMark-ETP, AUGUSTUS and TSEBRA. bioRxiv.

54. Tegenfeldt, F., Kuznetsov, D., Manni, M., Berkeley, M., Zdobnov, E. M. and Kriventseva, E. V. 2025, OrthoDB and BUSCO update: annotation of orthologs with wider sampling of genomes. Nucleic Acids Res, 53, D516–D522.

55. Hart, A. J., Ginzburg, S., Xu, M. S., et al. 2020, EnTAP: Bringing faster and smarter functional annotation to non-model eukaryotic transcriptomes. Mol Ecol Resour, 20, 591–604.

56. Buchfink, B., Reuter, K. and Drost, H. G. 2021, Sensitive protein alignments at tree-of-life scale using DIAMOND. Nat Methods, 18, 366–368.

57. Ou, S., Su, W., Liao, Y., et al. 2019, Benchmarking transposable element annotation methods for creation of a streamlined, comprehensive pipeline. Genome Biol, 20, 275.

58. Contreras-Moreira, B., Filippi, C. V., Naamati, G., Garcia Giron, C., Allen, J. E. and Flicek, P. 2021, K-mer counting and curated libraries drive efficient annotation of repeats in plant genomes. Plant Genome, 14, e20143.

59. Nussbaumer, T., Martis, M. M., Roessner, S. K., et al. 2013, MIPS PlantsDB: a database framework for comparative plant genome research. Nucleic Acids Res, 41, D1144–1151.

60. Amselem, J., Cornut, G., Choisne, N., et al. 2019, RepetDB: a unified resource for transposable element references. Mob DNA, 10, 6.

61. Wicker, T., Matthews, D. E. and Keller, B. 2002, TREP:a database for Triticeae repetitive elements. Trends in Plant Science, 7, 561–562.

62. Chan, P. P., Lin, B. Y., Mak, A. J. and Lowe, T. M. 2021, tRNAscan-SE 2.0: improved detection and functional classification of transfer RNA genes. Nucleic Acids Res, 49, 9077–9096.

63. Lagesen, K., Hallin, P., Rodland, E. A., Staerfeldt, H. H., Rognes, T. and Ussery, D. W. 2007, RNAmmer: consistent and rapid annotation of ribosomal RNA genes. Nucleic Acids Res, 35, 3100–3108.

64. Nawrocki, E. P. and Eddy, S. R. 2013, Infernal 1.1: 100-fold faster RNA homology searches. Bioinformatics, 29, 2933–2935.

65. Kalvari, I., Nawrocki, E. P., Ontiveros-Palacios, N., et al. 2021, Rfam 14: expanded coverage of metagenomic, viral and microRNA families. Nucleic Acids Res, 49, D192–D200.

66. Kozomara, A., Birgaoanu, M. and Griffiths-Jones, S. 2019, miRBase: from microRNA sequences to function. Nucleic Acids Res, 47, D155–D162.

67. Krzywinski, M., Schein, J., Birol, I., et al. 2009, Circos: an information aesthetic for comparative genomics. Genome Res, 19, 1639–1645.

68. Cabanettes, F. and Klopp, C. 2018, D-GENIES: dot plot large genomes in an interactive, efficient and simple way. PeerJ, 6, e4958.

69. Goel, M., Sun, H., Jiao, W. B. and Schneeberger, K. 2019, SyRI: finding genomic rearrangements and local sequence differences from whole-genome assemblies. Genome Biol, 20, 277.

70. Hao, Z., Lv, D., Ge, Y., et al. 2020, RIdeogram: drawing SVG graphics to visualize and map genome-wide data on the idiograms. PeerJ Comput Sci, 6, e251.

71. Jain, C., Rodriguez, R. L., Phillippy, A. M., Konstantinidis, K. T. and Aluru, S. 2018, High throughput ANI analysis of 90K prokaryotic genomes reveals clear species boundaries. Nat Commun, 9, 5114.

72. Wang, Y., Tang, H., Debarry, J. D., et al. 2012, MCScanX: a toolkit for detection and evolutionary analysis of gene synteny and collinearity. Nucleic Acids Res, 40, e49.

73. Emms, D. M. and Kelly, S. 2019, OrthoFinder: phylogenetic orthology inference for comparative genomics. Genome Biol, 20, 238.

74. Emms, D. M. and Kelly, S. 2015, OrthoFinder: solving fundamental biases in whole genome comparisons dramatically improves orthogroup inference accuracy. Genome Biol, 16, 157.

75. Cazaux, B., Catalan, J., Veyrunes, F., Douzery, E. J. and Britton-Davidian, J. 2011, Are ribosomal DNA clusters rearrangement hotspots? A case study in the genus Mus (Rodentia, Muridae). BMC Evolutionary Biology, 11, 124.

76. Nakajima, R. T., Cabral-de-Mello, D. C., Valente, G. T., Venere, P. C. and Martins, C. 2012, Evolutionary dynamics of rRNA gene clusters in cichlid fish. BMC Evolutionary Biology, 12, 198.

77. Wang, W., Zhang, X., Garcia, S., Leitch, A. R. and Kovarik, A. 2023, Intragenomic rDNA variation - the product of concerted evolution, mutation, or something in between? Heredity, 131, 179–188.

78. Roa, F. and Guerra, M. 2012, Distribution of 45S rDNA sites in chromosomes of plants: Structural and evolutionary implications. BMC Evolutionary Biology, 12, 225.

79. Garcia, S., Kovarik, A., Leitch, A. R. and Garnatje, T. 2017, Cytogenetic features of rRNA genes across land plants: analysis of the Plant rDNA database. Plant J, 89, 1020–1030.

80. Stevison, L. S., Hoehn, K. B. and Noor, M. A. F. 2011, Effects of inversions on within- and between-species recombination and divergence. Genome Biology and Evolution, 3, 830–841.

81. Huang, K. and Rieseberg, L. H. 2020, Frequency, origins, and evolutionary role of chromosomal inversions in plants. Frontiers in Plant Science, 11, 296.

82. Villoutreix, R., Ayala, D., Joron, M., Gompert, Z., Feder, J. L. and Nosil, P. 2021, Inversion breakpoints and the evolution of supergenes. Molecular Ecology, 30, 2738–2755.

83. Bock, D. G., Cai, Z., Elphinstone, C., et al. 2023, Genomics of plant speciation. Plant Communications, 4, 100599.

84. Palmer, J. D. and Shields, C. R. 1984, Tripartite structure of the Brassica campestris mitochondrial genome. Nature, 307, 437–440.

85. Kubo, T. and Newton, K. J. 2008, Angiosperm mitochondrial genomes and mutations. Mitochondrion, 8, 5–14.

86. Gualberto, J. M., Mileshina, D., Wallet, C., Niazi, A. K., Weber-Lot, F. and Dietrich, A. 2014, The plant mitochondrial genome: dynamics and maintenance. Biochimie, 100, 107–120.

87. Bendich, A. J. 1993, Reaching for the ring: the study of mitochondrial genome structure. Current Genetics, 24, 279–290.

88. Bendich, A. J. 2010, Mitochondrial DNA, chloroplast DNA and the origins of development in eukaryotic organisms. Biology Direct, 5, 42.

89. Cheng, N., Lo, Y.-S., Ansari, M. I., et al. 2017, Correlation between mtDNA complexity and mtDNA replication mode in developing cotyledon mitochondria during mung bean seed germination. New Phytologist, 213, 751–763.

