## Supplementary Fig. for "Chromosome-scale genomes of two wild flowering cherrys (*Cerasus itosakura* and *C. jamasakura*) provide insights into structural evolution in *Prunus*"

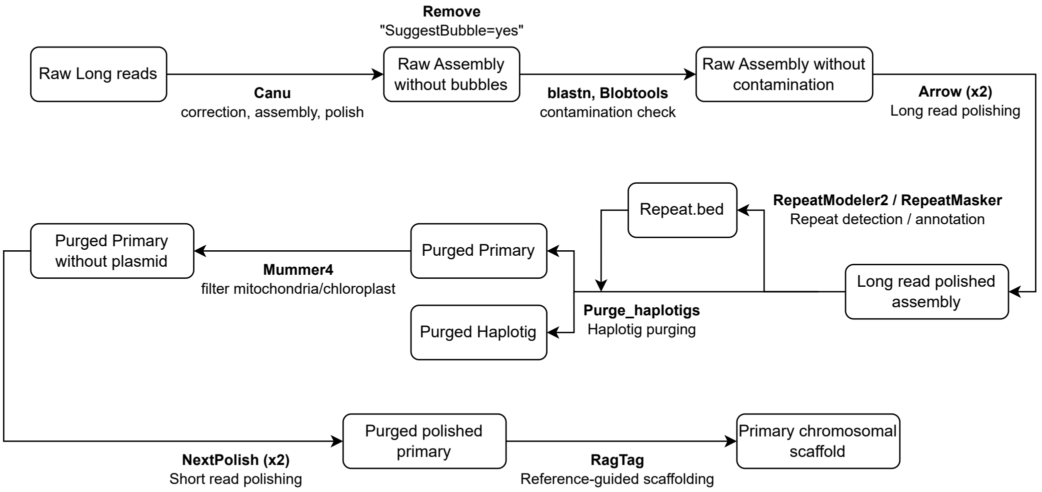


**Supplementary Figure 1: Overall workflow of the genome assembly**

This figure illustrates the overall genome assembly workflow up to the reference-guided scaffolding stage, with the arrows indicating the tools used at each step.

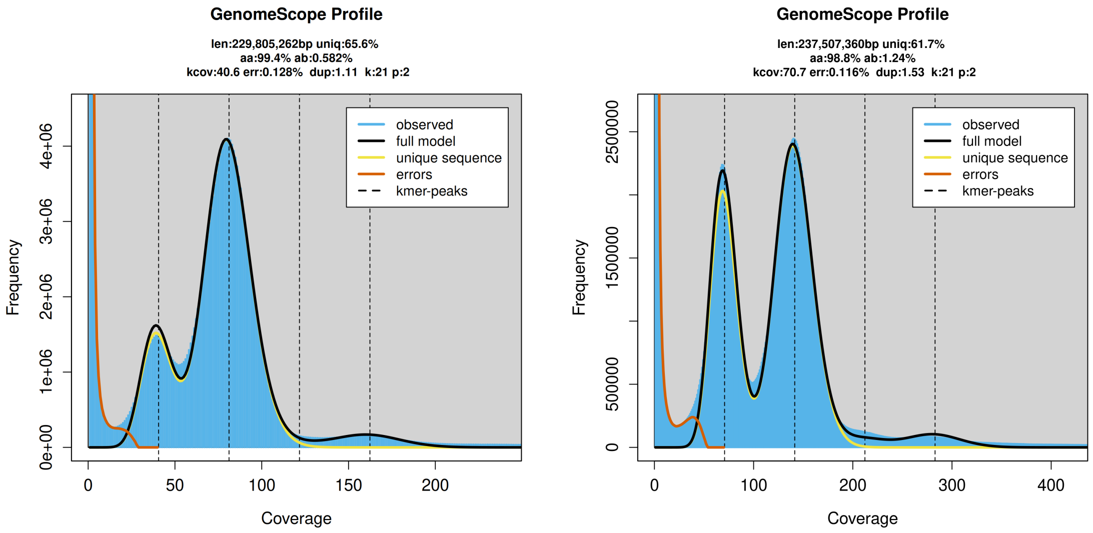


**Supplementary Figure 2: k-mer frequency distributions**

Genome size and heterozygosity for the two samples analyzed in this study were estimated from k-mer frequency distributions generated from short-read data. The left panel shows the results for *C. itosakura* and the right panel for *C. jamasakura*. Assuming a diploid genome, the haploid genome size was estimated to be 229,805,262 bp for *C. itosakura* and 237,507,360 bp for *C. jamasakura*. The estimated heterozygosity was 0.58% and 1.24%, respectively, indicating a higher level of heterozygosity in *C. jamasakura*.

*Cerasus itosakura*

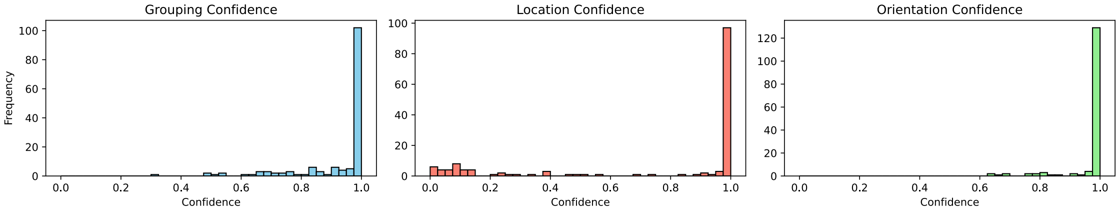


*Cerasus jamasakura*

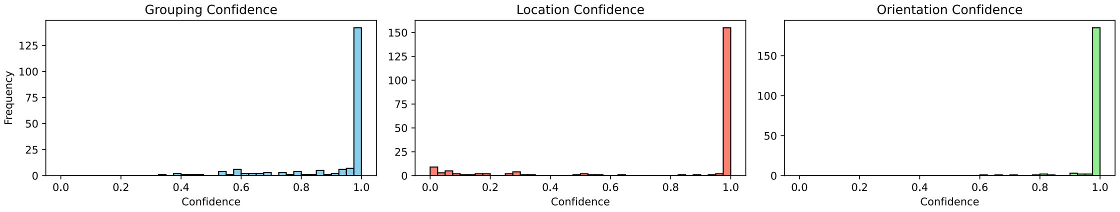


**Supplementary Figure 3: Distribution of confidence scores in RagTag scaffolding for *C. itosakura* and *C. jamasakura***

Histograms show the distribution of confidence scores from homology-based scaffolding using the complete genome of *Cerasus speciosa* as a reference. The distributions are shown for *Cerasus itosakura*, and *Cerasus jamasakur*a.

The grouping confidence score reflects the reliability of assigning a contig to a particular reference sequence (chromosome or scaffold). The location confidence score indicates the certainty of placing the contig at a specific position within the reference. The orientation confidence score represents the confidence in determining the correct direction of the contig (forward or reverse complement). In all cases, scores closer to 1 indicate higher confidence.

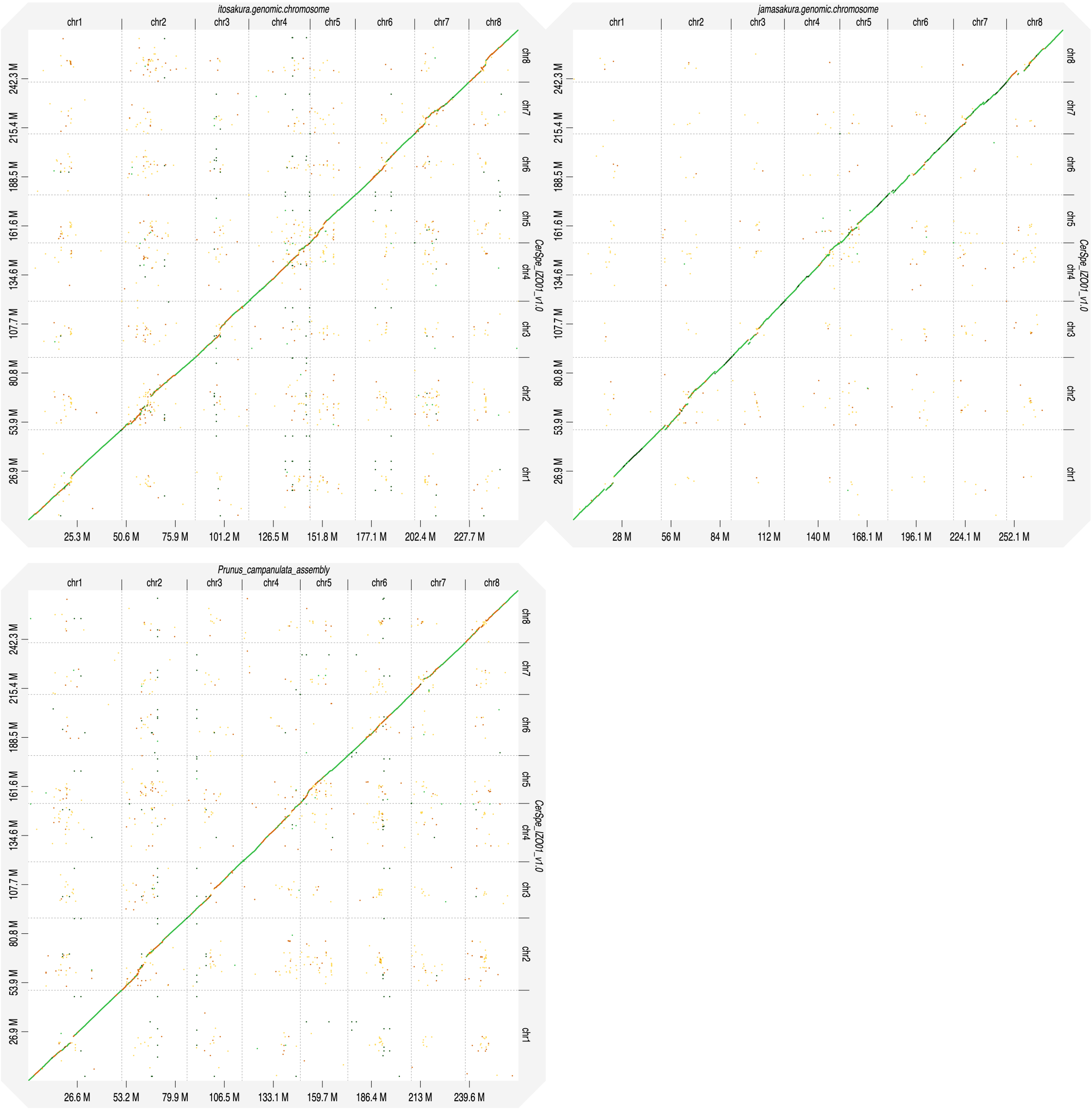


**Supplementary Figure 4: Comparative genome structure analysis using dot plots**

Dot plots generated using *C. speciosa* as the reference. The upper left panel shows the comparison with *C. itosakura*, the upper right with *C. jamasakura*, and the lower left with *C. campanulata*. The y-axis represents the *C. speciosa* genome, and the x-axis represents the genome of the comparison species. Darker green dots indicate higher sequence similarity, whereas darker orange dots indicate lower similarity. Overall, *C. itosakura*, *C. jamasakura*, and *C. campanulata* show high structural and sequence similarity to *C. speciosa*, with *C. jamasakura* exhibiting the highest sequence similarity.

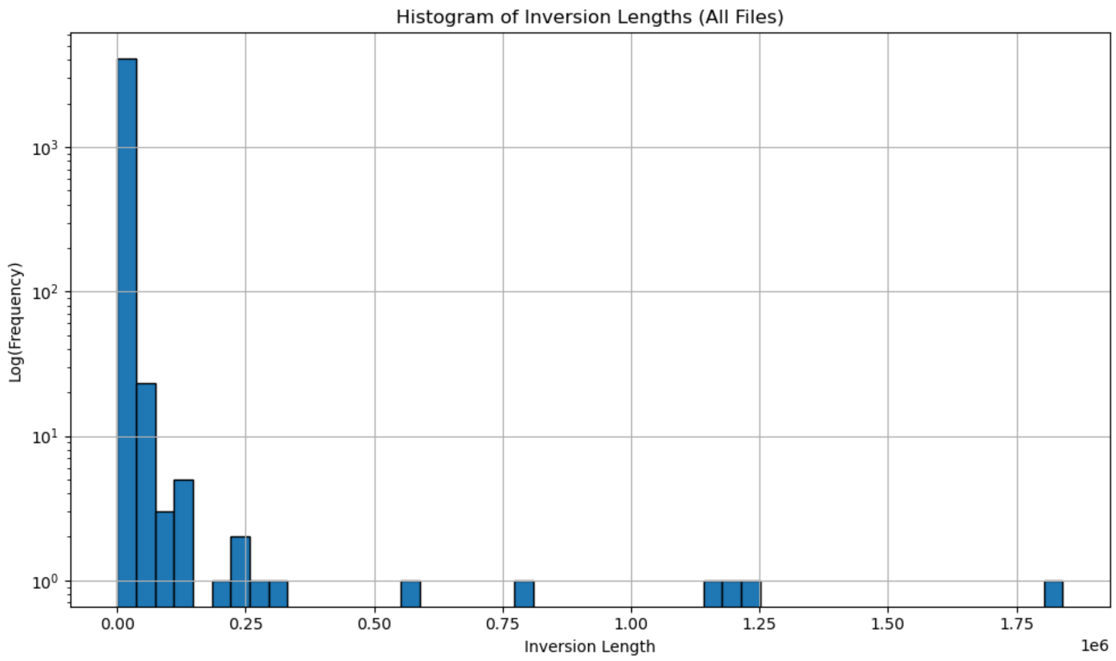


**Supplementary Figure 5: Histogram of chromosome inversion lengths**

Histogram comparing chromosome inversion lengths when the *C. speciosa* genome was used as the reference and the genomes of *C. itosakura*, *C. jamasakura*, and *C. campanulata* were used for comparison. The x-axis represents inversion length (bp), and the y-axis shows the frequency on a logarithmic scale.

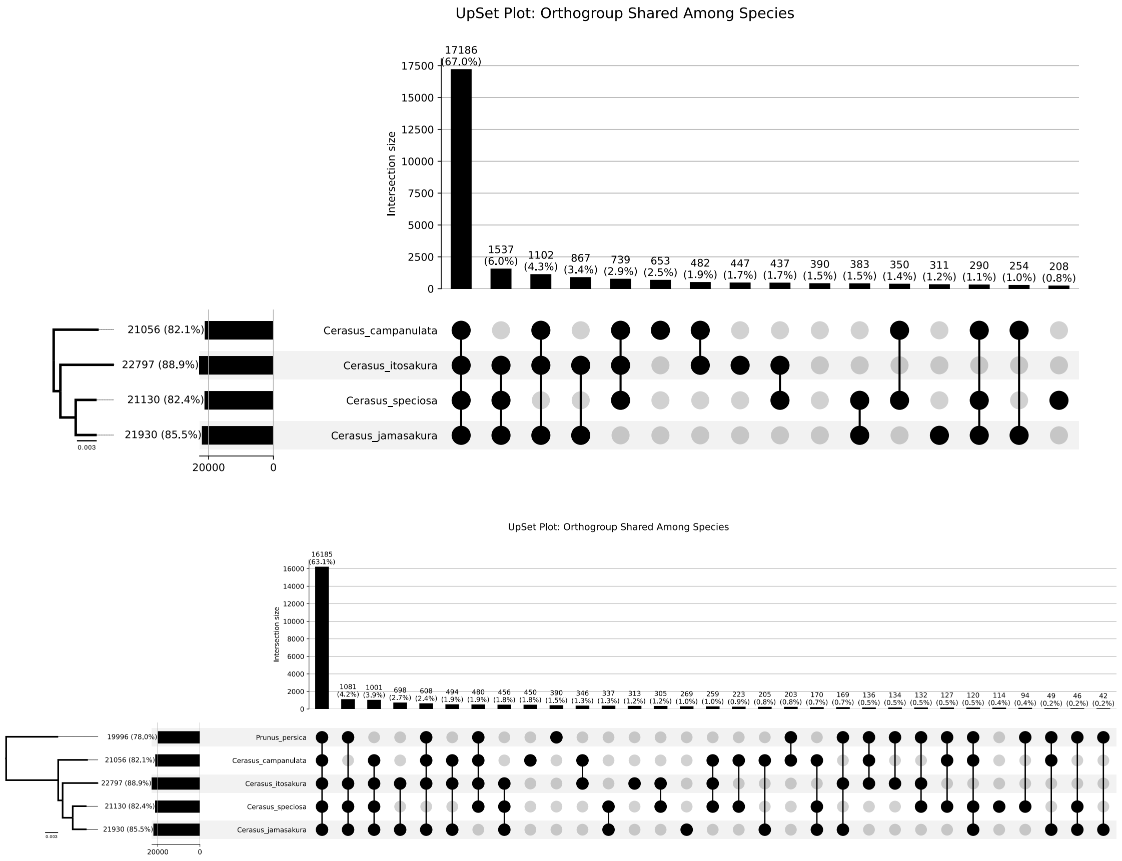


**Supplementary Figure 6: UpSet plot comparing orthologous gene sets**

(Top) UpSet plot showing the distribution of orthologous gene sets identified among *C. itosakura*, *C. jamasakura*, *C. speciosa*, and *C. campanulata*. The phylogenetic tree on the left illustrates the inferred relationships among the four species. Horizontal bars represent the number of genes in each shared or unique orthologous set, and connected dots below indicate the species included in each set.

(Bottom) UpSet plot generated using the same approach, but including *P. persica* as an outgroup. The addition of *P. persica* reduced the proportion of genes shared across all Cerasus species from 67.0% to 63.1%.

*Cerasus* × *yedoensis* (*Cerasus itosakura* haplotype)

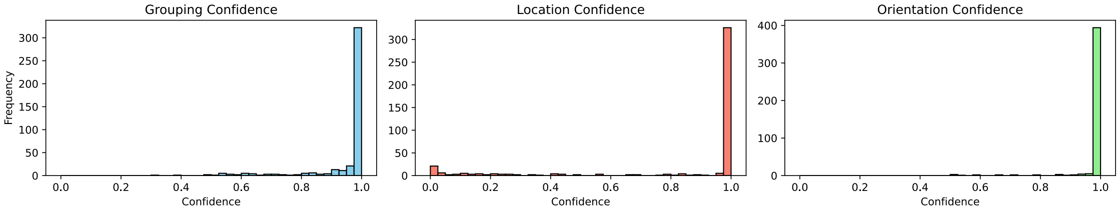


*Cerasus* × *yedoensis* (*Cerasus speciosa* haplotype)

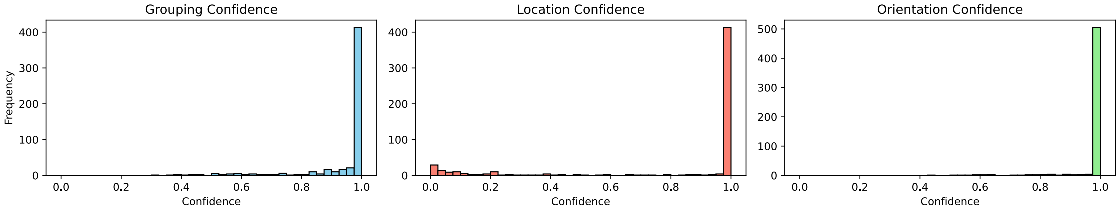


**Supplementary Figure 7: Distribution of Confidence Scores in RagTag Scaffolding for *C.* ×*yedoensis***

Histograms show the distribution of confidence scores from homology-based scaffolding. For the *C.* ×*yedoensis* genome, the *C. speciosa* haplotype was scaffolded using the complete genome sequence of *C. speciosa* as the reference, whereas the *C. itosakura* haplotype was scaffolded using the chromosome-level genome assembly of *C. itosakura* generated in this study. The distributions are shown for *C.* ×*yedoensis*, encompassing both the *C. itosakura* and *C. speciosa* haplotypes.

The grouping confidence score reflects the reliability of assigning a contig to a particular reference sequence (chromosome or scaffold). The location confidence score indicates the certainty of placing the contig at a specific position within the reference. The orientation confidence score represents the confidence in determining the correct direction of the contig (forward or reverse complement). In all cases, scores closer to 1 indicate higher confidence.

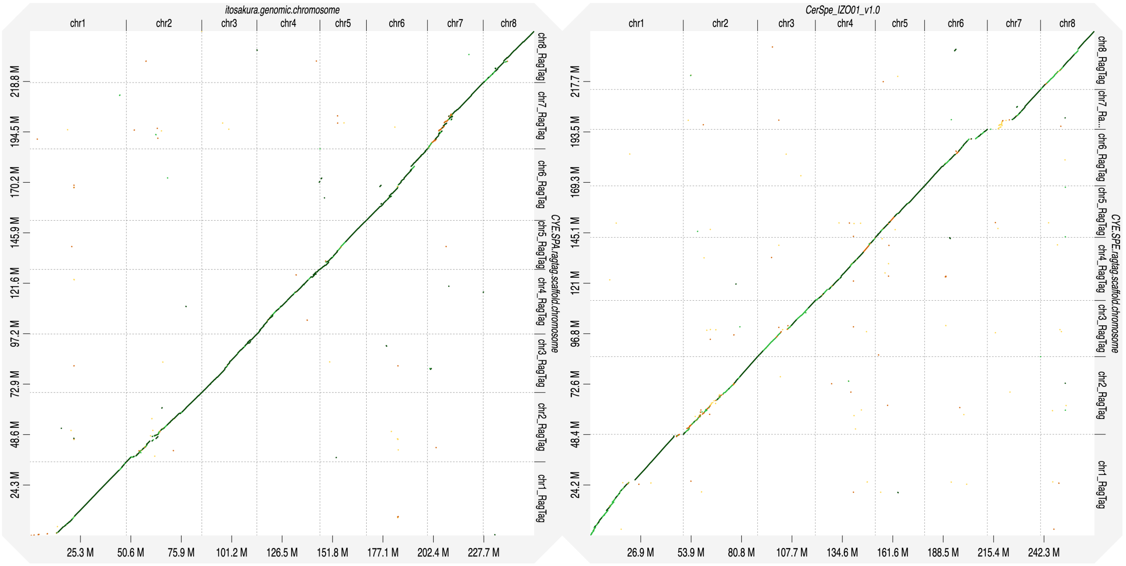

**Supplementary Figure 8: Comparative genome structure analysis using dot plots for *C. ×yedoensis***

The left panel shows the comparison between the reconstructed *C. ×yedoensis C. itosakura* haplotype and the newly assembled *C. itosakura* genome, and the right panel shows the comparison between the reconstructed *C. ×yedoensis C. speciosa* haplotype and the *C. speciosa* genome. Darker green dots indicate higher sequence similarity, whereas darker orange dots indicate lower similarity. Both haplotypes of *C. ×yedoensis* exhibit very high sequence similarity to their respective reference genomes. However, certain regions of the *C. ×yedoensis* genome show a loss of sequence information.

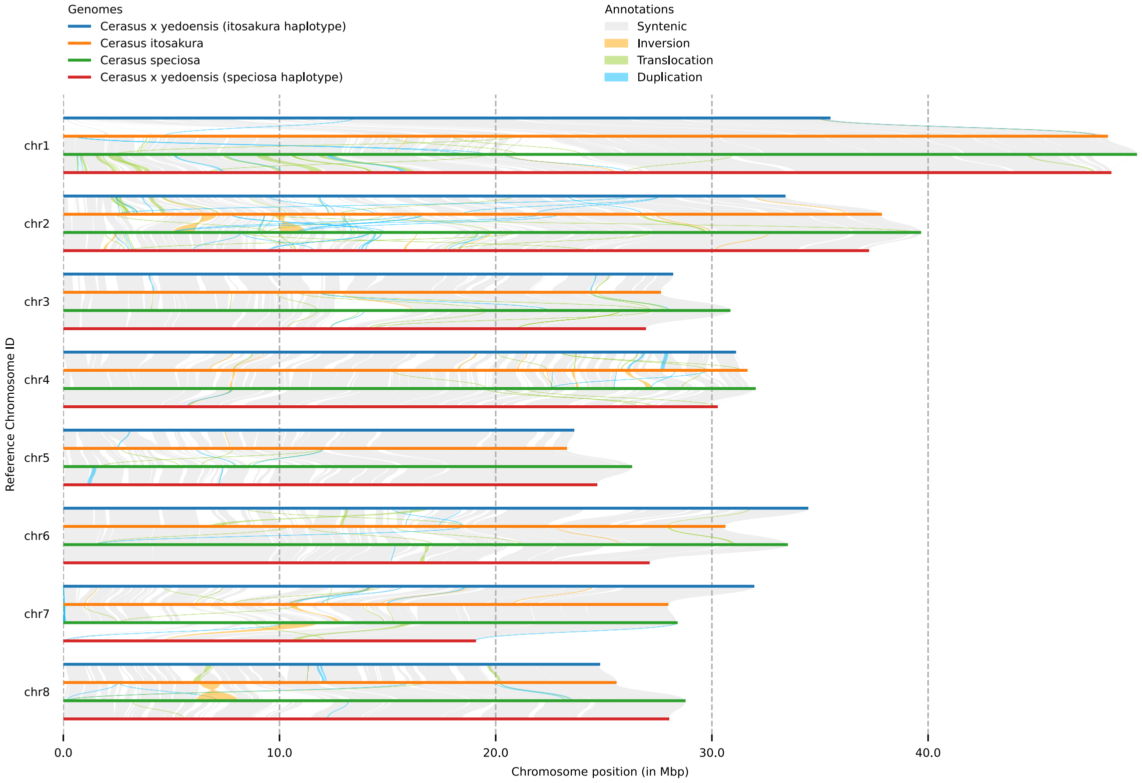


**Supplementary Figure 9: Chromosome-scale synteny between each haplotype of *C.* ×*yedoensis* and its respective wild progenitor species.**

Comparative chromosome alignments of *C.* ×*yedoensis* (both *C. itosakura* and *C. speciosa* haplotypes), *C. itosakura*, and *C. speciosa*. Alignments are colored according to structural relationships, including conserved synteny, inversions, translocations, and duplications.
